# The Type VI secretion effector TpeX expands the pore-forming virulence arsenal of *Pseudomonas aeruginosa*

**DOI:** 10.64898/2026.08.24.746647

**Authors:** Chantal Soscia, Sébastien Reig, Delphine Lefebvre, Maximilien Rouzaud, Laura Schmitt, Bérengère Ize, Gael Brasseur, Sophie Bleves

## Abstract

The type VI secretion system (T6SS) is a major weapon used by *Pseudomonas aeruginosa* to antagonize competing bacteria through the delivery of a diverse repertoire of toxic effectors. Although several *P. aeruginosa* T6SS effectors target bacterial membranes, the mechanisms underlying membrane disruption remain poorly characterized. Here, we study TpeX (PA5265), an accessory T6SS effector from *P. aeruginosa* PAO1 with structural similarity to the VasX pore-forming effector of *Vibrio cholerae*. We show that membrane-targeted TpeX exerts a bactericidal activity in *Escherichia coli*, resulting in dissipation of the membrane potential and loss of membrane integrity. The TpeX C-terminal region containing the predicted colicin-like transmembrane domain is sufficient to confer toxicity, although with reduced activity, supporting its role as the membrane-disrupting module. TpeX also oligomerizes upon membrane targeting, forming at least dimers, and structural modelling predicts a membrane-embedded pore compatible with the observed permeabilization phenotype. We further identify TpiX (PA5264), the protein encoded by the downstream gene, as the cognate immunity protein, which partially protects cells from TpeX toxicity and interacts with TpeX. Finally, AlphaFold 3 modelling predicts an interaction between TpeX and the HcpB-VgrG6 T6SS spike, suggesting a possible mechanism for effector recruitment and delivery. Together, our results identify TpeX as a bactericidal, colicin-like pore-forming T6SS effector whose membrane activity is controlled by a cognate immunity protein, thereby expanding the repertoire of membrane-targeting weapons used by *P. aeruginosa* in interbacterial competition.

## INTRODUCTION

Antimicrobial resistance is a multifactorial phenomenon recognized as a public health threat by the World Health Organization that could cause a disastrous crisis in a near future^1^. The combination of bactericidal properties of antibiotics on one side and well-known evolutionary pressure mechanisms on the other side favors the appearance of resistant bacteria^2^. Consequently, some *Pseudomonas aeruginosa* isolates have become resistant to all available treatments, highlighting the current unmet medical need in the field. In this context, bacterial virulence targeting strategy is a promising paradigm, with the potential through disarming and non-killing modes of action to attenuate the selective pressure responsible of drug resistance or to promote the selection of less virulent strains ^2–4^. Unfortunately, to date no anti-virulent agent has been introduced in routine clinical practice against *P. aeruginosa* underscoring the need to identify and characterize novel and relevant anti-virulence targets to reimagine patients care^5,6^.

*P. aeruginosa* is an opportunistic pathogen characterized by a large arsenal of virulence factors including motility, adherence, the ability to form a biofilm under certain conditions and to secrete many toxins. These toxins also called effectors are released in the surrounding environment or injected directly into host cells or into other competing bacteria^7^. Discovered in 2006 in *Vibrio cholerae* and in *P. aeruginosa*^8,9^ and widely conserved among Gram-negative bacteria, the type VI secretion system (T6SS) is a contractile nanomachine anchored in the bacterial envelope allowing the secretion of the associated type VI effectors. Contraction of the T6SS sheath can propel an effector-loaded arrow into bacterial and/or eukaryotic target cells. The arrow is a nanotube composed of a stack of Hcp proteins, topped with a perforating spike (an assembly of VgrG and PAAR proteins). Effectors can either be loaded inside the Hcp tube or associated with the spike complex (cargo effectors) or fused to components of the secretion machinery (evolved VgrG, PAAR, or Hcp)^10^. Some effectors need a cytoplasmic adaptor, also called chaperone, to be targeted to the T6SS machinery. *P. aeruginosa* T6SS is encoded at three distinct *loci*, H1-T6SS involved only in antibacterial activity, H2-T6SS and H3-T6SS targeting bacterial or eukaryotic cells and thus expressing trans-kingdom activity effectors able to target both types of cells^11–13^. In addition to these core elements, orphan *vgrG loci* scattered around the chromosome have also been found to belong to the T6SS^14–16^. They are considered predictive of T6SS effector candidates because genes encoding effectors and those involved in their secretion are found associated downstream of them^14,17–19^.

*P. aeruginosa* T6SS delivers a cocktail of effectors, 20 for the PAO1 strain, with broad range of targets and various activities^16^: (*i*) cell wall peptidoglycan (Tse1: amidase^20^; Tse3: muramidase^21^), (*ii*) membrane (Tle1, Tle3, Tle4 and Tle5a and Tle5b: phospholipases^22^), (*iii*) inner membrane (Tse4, Tse5: pore-forming activity^23,24^), (*iv*) DNA (Tse7: nuclease^25^), (*v*) NAD(P)^+^ (Tse6: hydrolase^26^), (*vi*) protein (metallopeptidase: VgrG2b^13^), (vii) transamidosome (Tse8: protein synthesis^27^), (viii) essential metabolic pathways (Tas1: depletion of ATP^28^) and (ix) metal acquisition system (TseF: iron uptake^29^). With these characteristics, the *P. aeruginosa* T6SS represents a key virulence factor actively killing or inhibiting the growth of surrounding competitor bacteria^30^, manipulating eukaryotic host cells^31^, and giving a competitive advantage in the dynamic ecosystem to access limited resources^29^. Consequently, targeting T6SS macromolecular assembly or its secreted effectors would interfere host colonization or escape from the immune system and could be an innovative and impactful strategy against *P. aeruginosa* infections. Many *P. aeruginosa* T6SS effectors mode of action remains uncharacterized to date and need to be elucidated.

A recent genomic analysis of the diversity of T6SS effectors among nearly 2,000 available *P. aeruginosa* genome sequences revealed core and accessory effectors of the three T6SS, classified according to their occurrence^16^. The orphan *vgrG6* island belongs to the set of accessory effectors of the H2-T6SS. Interestingly, this locus shows especially rare diversity among strains and is subject to recombination with 3 mutually exclusive effector genes, or loss of the effector-coding gene. Indeed, the PAO1 strain is representative of strains carrying *tpsE1b* (PA5265), while the PA14 strain carries *tpsE1c* (*ptx2*, PA14_69520) and the PALA52 strain is even more unique because the *vgrG6* island is not at the same localization on the chromosome and possesses another effector gene *tpsE1a* (PALA52_0091)^15,16^. Ptx2 and PA5265 are homologous T6SS effectors that have undergone substantial sequence diversification, particularly in their C-terminal domains. In this study, we focused on the product of the PA5265 gene of the PAO1 strain. PA5265, as Ptx2 of the PA14 strain^32,33^, is highly similar to the VasX effector of *V. cholerae*. VasX is known as the first T6SS antibacterial and anti-eukaryotic pore-forming toxin^34–36^. VasX dissipates the cell membrane potential, permeabilizes target cells, and binds phospholipids, indicating that it acts as a pore-forming colicin^37–40^.

In this study, we report the characterization of PA5265 of *P. aeruginosa* PAO1. Firstly, a strong predicted structural homology of PA5265 (TpeX) with VasX of *V. cholerae*^34–36^ has oriented our experiments toward a membrane targeting function. Indeed, TpeX targeted to the inner membrane, can inhibit bacterial cells growth in heterologous conditions. This toxicity is counteracted by a cognate immunity protein TpiX. Using DiOC2(3) and propidium iodide dyes fluorescence measured with flow cytometry, we demonstrate that TpeX induced toxicity was corelated to membrane potential dissipation and membrane integrity disruption. Oligomerization of TpeX and inner membrane pore formation could be responsible for the bactericidal effect of this T6SS effector of *P. aeruginosa*. We used AlphaFold to predict the complex of TpeX with structural components of the T6SS machinery. The resulting model predicts with confidence the interaction of the TpeX and VgrG6 tip of an HcpB tube, thus strengthening our hypothesis.

## RESULTS

### TpeX (PA5265) is a putative T6SS dependent pore-forming effector

The orphan *vgrG6 locus* of *P. aeruginosa* PAO1 contains two other genes of unknown function, PA5264 and PA5265 (Fig. 1A). A σ70 promoter is predicted by BProm upstream of the *hcpB* gene and two more upstream of the PA5264 gene (Pseudomonas.com). Given this genetic organization, PA5264 and PA5265 genes could be predicted to encode an immunity-effector pair dependent on a HcpB-VgrG6 arrow for delivery. PA5265 encodes a hypothetical protein of 119.2 kDa predicted to be localized in the cytoplasmic membrane (Pseudomonas.com). Analysis of its sequence by DAS and TMHMM V2.0 predicts a large N-terminal domain in the periplasm followed by 3 predicted hydrophobic transmembrane helices (TMHs) and a C-terminal domain in the cytoplasm (Fig. S1A). This particular effector topology is reminiscent of that of VasX (VCA0020) topology of *V. cholerae.* Since these two proteins share the same topology, size and gene cluster organization (Fig. 1B), we hypothesize that PA5265 could be a VasX-like T6SS effector.

**Fig. 1.**
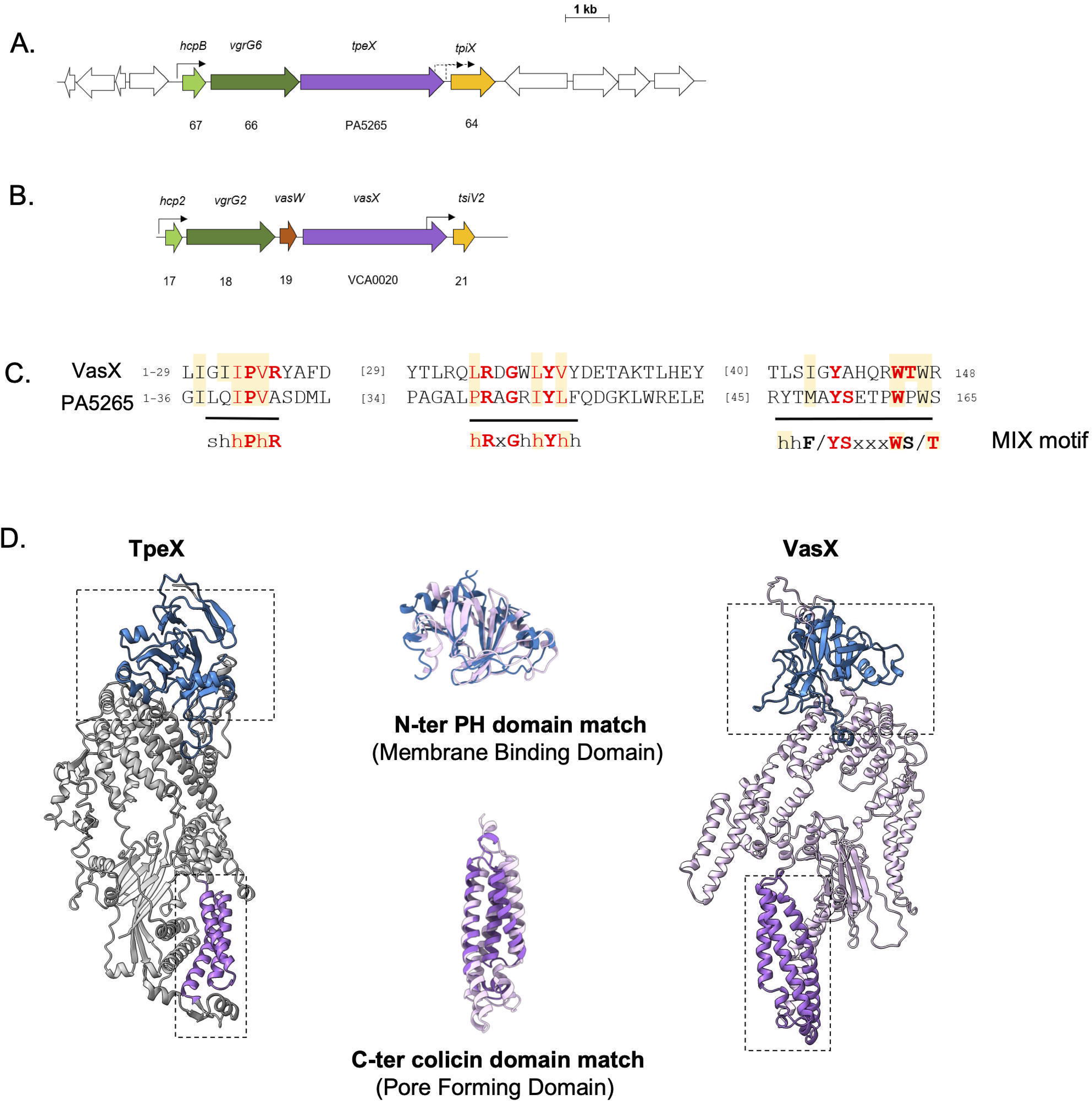
*PA5265* belongs to an orphan *hcp/vgrG* pathogenicity locus and codes a putative T6SS dependent pore­forming effector. (A) Schematic representation of *P. aeruginosa* PAO1 PA5264-5267 genes organization (Pseudomonas.com). White arrows correspond to non T6SS related genes. Internal a70 promotors in dotted lines were predicted using BPROM. **(B)** Schematic representation of *V. cholerae* V52 *VCA0017-0021* genes organization^36^. **(C)** Alignment of MIX (Marker for type sIX secretion system effectors) motif at the N-terminus of VasX and PA5265. MIX^41^ is composed of a widely conserved central sequence, hRxGhhYhh and by two less conserved sequences shhPhR and hhF/YSxxxWS/T. MIX motif conserved residues are highlighted in red. The figures indicate the residues not represented in the sequence alignment(D) Putative structural models homologies between VasX and PA5265 (TpeX) predicted by AlphaFold 2. Colicin domains with hydrophobic transmembrane helices are represented in purple; pleckstrin homology (PH) domains with anti-parallel beta sheets, followed by an amphipathic helix highlighted in blue and and outlined with a dotted line. N-ter, N-terminal end; C-ter, C-terminal end.

With 3 TMHs in a colicin domain, VasX is known to share structural homology with pore-forming colicin^36^. *In silico* analysis of the PA5265 C-terminus sequence using homology prediction programs revealed a putative colicin A domain [amino acids 742-840] (HHpred 54% probability; Phyre2 66% confidence and 16% identity). Among other *in silico* informative elements, a class 1 MIX motif (<u>M</u>arker for type s<u>IX</u> effectors) characterized in the N-terminal sequence of VasX^18,41^ was also found in the N-terminal sequence of PA5265 (Fig. 1C). Many class 1 MIX proteins contain predicted TMHs in C-terminus suggesting they could act as pore-forming toxins (PFTs)^41^.

As the structure of PA5265 remains unresolved, we relied on the structure predictions available in the AlphaFold 2 database (AFDB) and searched for structural homologies using the Foldseek search server. Interestingly, the first hit for PA5265 against AFDB-SWISSPROT was *V. cholerae* O1 VasX (100% probability; E-value 9.39e^-13^). Because membrane localization is not considered in the prediction, the conformation predicted by AlphaFold 2 is very probably not the active form of the toxin but rather corresponds to its soluble state. Indeed, PFTs are known to undergo structural transformations from soluble, inactive monomers to active, multimeric transmembrane pores that insert into the membranes of target cells^42^. A domain by domain structural comparison of the two proteins was nonetheless possible: it showed an overlap of the C-terminal colicin domains (Fig. 1D) and another overlap in the N-terminal region (Fig. 1D), suggesting that PA5265 also possesses a Pleckstrin Homology (PH) domain [amino acids 60-296], a domain crucial for VasX-mediated virulence and implicated in membrane binding to phosphatidylinositol lipids in eukaryotic proteins^43,36^. On the basis of the orphan T6SS gene cluster organization and the structural elements predicted to be homologous to VasX, we considered PA5265 as a putative T6SS colicin-like PFT and named it TpeX for <u>T</u>6SS dependent <u>p</u>ore-forming <u>e</u>ffector X according to the proposed nomenclature for Ssp6/Tpe1 of *Serratia marcescens*^44^.

Antibacterial effector genes are usually encoded with a cognate immunity gene preventing T6SS-dependent killing by neighboring cells and avoiding a self-toxic activity for effectors active in the cytoplasm^19^. The toxic effect of VasX is prevented by an immunity protein, TsiV2^35,36^, encoded downstream of *vasX* (Fig. 1B). We thus reasoned that PA5264, located downstream of *tpeX* might encode a TpeX immunity protein (Fig. 1A). Interestingly and in agreement with this proposed essential role of immunity protein, PA5264 was previously identified as an essential gene in *P. aeruginosa* PAO1 by a shotgun antisense screening approach^45^. The presence of two predicted internal promoters upstream of PA5264, as is similarly observed upstream of *tsiV2*^36^ (Fig. 1A & 1B), is also in line with an immunity function for PA5264. Indeed, in Type II toxin/antitoxin modules, the gene encoding the toxin is generally located downstream of the antitoxin gene leading to an excess of antitoxin after translation. But in case of an inverse order, like for *tpeX* and *vasX*, the antitoxin gene is transcribed with the toxin gene but also separately from internal promoters when the toxin to antitoxin ratio becomes too high^46^. Immunity proteins are known to localize to the cellular compartment in which the cognate effector exerts its activity^47^. Like TpeX, PA5264 is predicted to localize in the inner membrane and could adopt a topology with 4 TMHs in its N-terminal region (Fig. S1B). This localization could allow PA5264 to counteract putative TpeX toxicity. PA5264 was then named TpiX for <u>T</u>6SS dependent <u>p</u>ore-forming effector immunity <u>X</u>.

### TpeX-mediated growth inhibition when targeted to the inner membrane

Consistent with a membrane targeting activity, pore-forming colicins and VasX must be targeted to and associated with the inner membrane of prey bacterium to be toxic^36^. To investigate whether this is also the case for TpeX, we have designed a heterologous toxicity assay *in E. coli*. This involves fusing a N-terminal signal sequence (SS) to direct the protein to the Sec translocon and to the periplasm. The *tpeX* sequence was cloned under a P_T7_ promoter in frame with the sequence encoding the PelB SS in the pET22b vector (called SS-TpeX), or in the pET-Duet (called TpeX) and with a His-tag sequence. Correct production and localization of these recombinant proteins in *E. coli* BL21(DE3) pLysS were verified by western blot after cell fractionation (Fig. 2A). Despite its 3 predicted TMHs, TpeX produced in *E. coli* remained cytoplasmic and was recovered only in the soluble fraction like the periplasmic control protein TolB, whereas SS-TpeX was associated both to the soluble and membrane fractions (Fig. 2A). Whereas the soluble TpeX does not impact *E. coli* growth (Fig. 2B, same growth as the strain carrying the empty vector), SS-TpeX inhibited *E. coli* growth on solid medium, demonstrating TpeX heterologous toxicity (Fig. 2B, 4 log difference between empty vector, TpeX and SS-TpeX). Next, we took advantage of this assay to delimit the toxic domain of TpeX. Sequences encoding the C-terminus of TpeX (from the first TMH to the end, amino acids 742-1089, Fig. S1A) or only the three TMHs (amino acids 742-840 corresponding to the colicin domain) were cloned downstream of the PelB SS. Both truncated proteins were toxic to *E. coli* albeit at a lower level (2 log difference) than the full-length SS-TpeX (Fig. 2B). As for VasX^36^, the colicin domain of TpeX is key to its toxic activity.

**Fig. 2.**
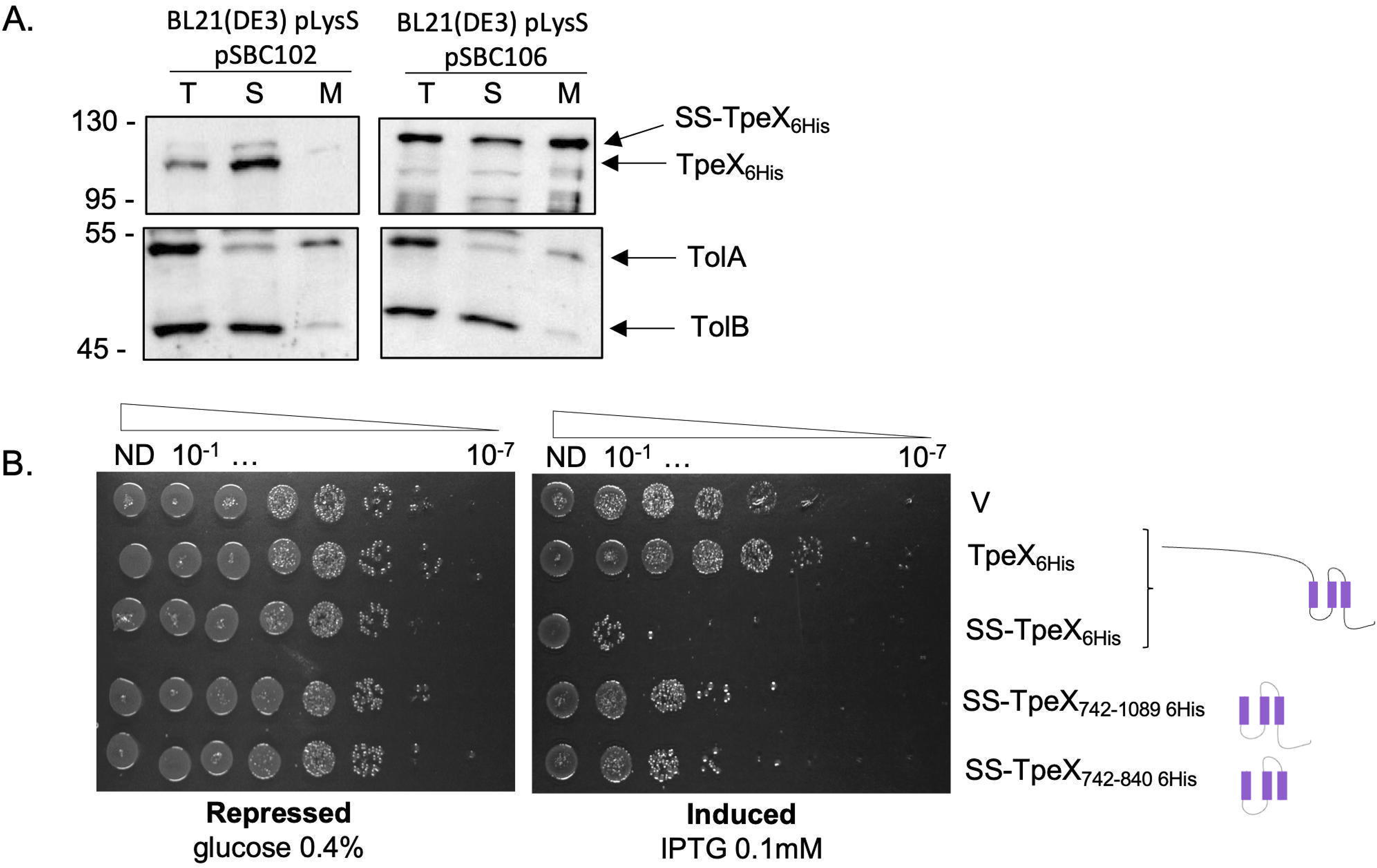
The heterologous toxicity of TpeX when targeted at the membrane. **(A)** Subcellular localization of TpeX_6H_i_S_ (from pSBC102) and SS-TpeX_6H_i_S_ (from pSBC106 yielding a fusion of TpeX with a Sec signal peptide) after *E. coli* fractionation and immunoblotting using antibodies directed against the His tag, TolB and TolA. TolB (55 kDa) and TolA (45 kDa) were used as periplasmic and inner membrane controls respectively. T: whole cell, S: soluble fraction (cytoplasm & periplasm), Mb: total membrane (inner and outer membranes). The position of the proteins and the molecular mass markers (in kDa) are indicated. **(B)** SS-TpeX_6H_i_S_ is toxic towards *E. coli.* Serial dilutions (from non-diluted to 10^-7^) of normalized cultures of *E. coli* producing the wild-type TpeX_6H_i_S_ (from pSB102) or targeted to the membrane, called SS-TpeX_6H_i_S_ (from pSB106), were spotted on LB agar plates supplemented with 0.4% glucose (left panel) or with 0.1 mM IPTG (right panel). Glucose and IPTG allow respectively repression and induction of the gene encoding the T7 RNA polymerase. Two truncated variants of SS-TpeX_6His_, produced from pSBC157 and pSBC158, were used as indicated, as well as the empty vector (V). The schemes on the right show the TpeX domains that are produced, the transmembrane segments are shown in purple.

### TpiX is the immunity protein of TpeX

To demonstrate an immunity role for TpiX, the gene was cloned in the pRSF-Duet vector to enable co-production with the toxin and, potentially, its neutralization. However, co-production of the putative immunity TpiX together with SS-TpeX could not counteract TpeX toxicity in *E. coli* and produced the same colony phenotype on solid medium as SS-TpeX produced alone (Fig. S2). Although this is a classic experiment, we recently encountered the same problem with Tli1a, the immunity protein from the Tle1 effector of *P. aeruginosa*^48^. Then, we developed another strategy common in the field of colicins^49,50^, a qualitative assay to assess the ability of bacteria, whether or not they produce TpiX, to grow after transformation by a vector encoding the toxic form of TpeX. To this end, *E. coli* was first transformed with the plasmid encoding TpiX or with the empty vector and these bacteria were then transformed with the plasmid encoding SS-TpeX. After the second transformation, half of the cells were spread onto a repressive medium (containing glucose 0.4%) and the other half onto an inducing medium (containing IPTG 0.01 mM). Very few transformants were recovered upon *E. coli* transformation with the plasmid encoding SS-TpeX on inducing medium whereas in a repressive medium, the transformation is highly effective (Fig. 3A, compare top left and top right). In line with this, we were able to achieve cloning *ss-tpeX* (pSBC106) by adding glucose to all cultures to repress expression. In contrast, a larger number of colonies were observed upon the transformation of *E. coli* already producing TpiX on inducing medium (Fig. 3A, bottom right). Therefore, the production of TpiX appears to provide partial protection against SS-TpeX-induced toxicity in *E. coli* under these experimental conditions.

**Fig. 3.**
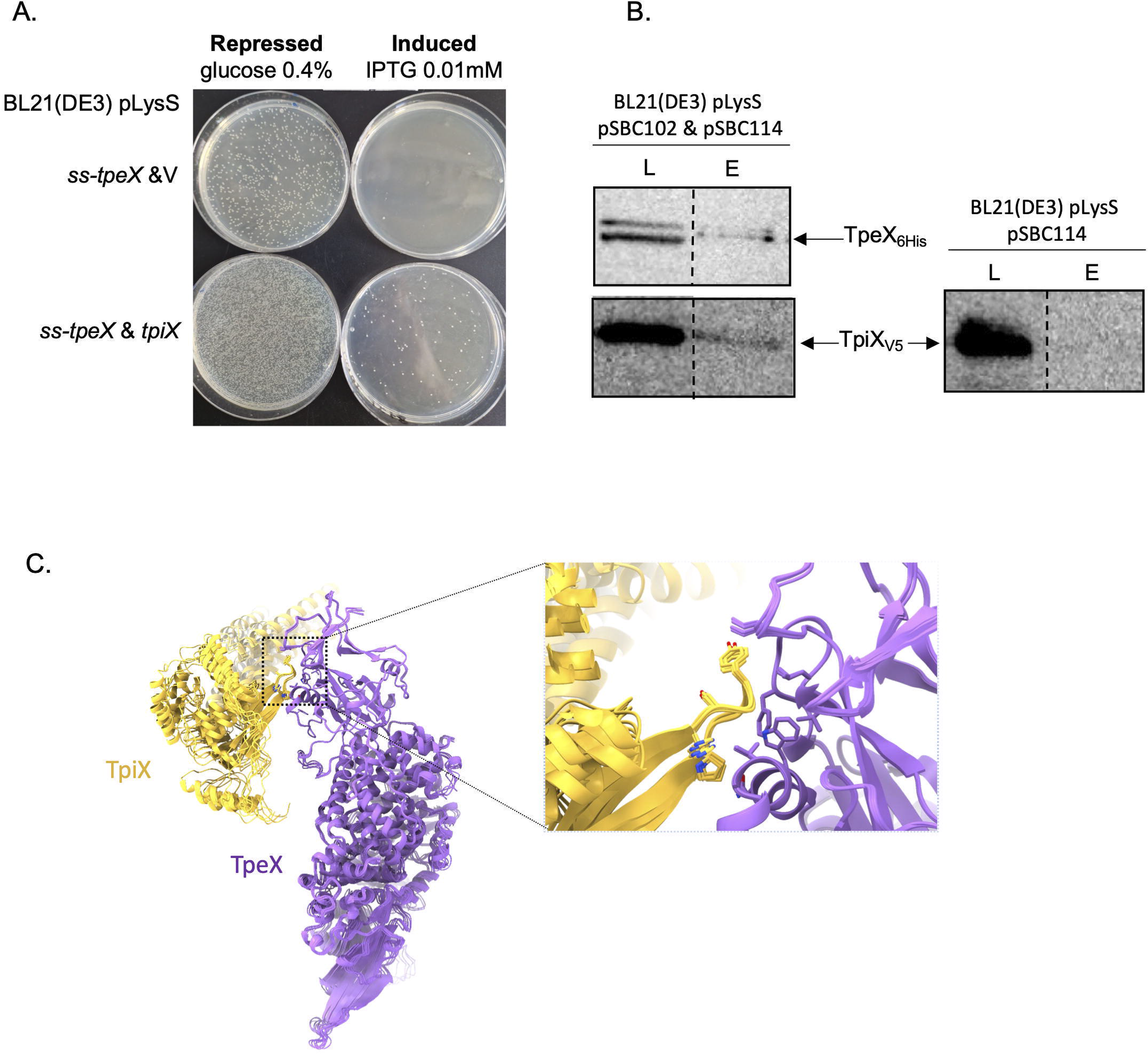
The TpiX immunity counteracts TpeX toxicity. **(A)** TpiX prevents SS-TpeX toxicity. Transformants selection on selective media containing glucose 0.4% (left panel) or IPTG 0.01 mM (right panel). Strains carrying *tpiX* (from pSBC114, lower panel) or not (pRSFDuet-1, upper panel) were transformed with the pSBC106 coding SS-TpeX. **(B)** Co-purification assays in batch with cobalt resin were done using BL21(DE3) pLysS to produce both TpeX_6His_ (from pSBC102) and TpiX_V5_ (from pSBC114) (left) or a TpiX_V5_ (right). The lysate (L) and eluted (E) fractions were collected and subjected to SDS-PAGE and Western blot analyses using anti-TpeX antibody (Upper) and anti-V5 antibody (Lower). **(C)** TpeX-TpiX interaction models generated by AlphaFold 3. In the inset, the overlapping of the 5 different models of the complex shows the conservation of the residues predicted to interact between the two proteins. In the models, TpiX is shown in yellow and TpeX in purple. The predicted distances between interacting residues are presented in Fig. S3).

To confirm an immunity role for TpiX, we studied the interaction between TpeX and TpiX by co-purification using affinity chromatography. We used the plasmid encoding the soluble and not toxic TpeX that allows the production of an His-tagged protein (from Fig. 2A) and the plasmid encoding TpiX tagged with a V5 tag (from Fig. 3A). The recombinant proteins were co-produced in *E. coli* BL21(DE3) pLysS. The bacterial lysate was loaded onto a cobalt matrix (see “Materials and Methods”), and TpeX_His6_ was eluted with imidazole. Despite the use of different production and purification conditions including various detergents, TpeX was very poorly purified (Fig. 3B). As showed in figure 3B, TpiX_V5_ was found in the eluted fraction only upon co-production with TpeX_His6_ (left panel). By contrast, when produced alone in *E. coli*, TpiX_V5_ was not retained by affinity chromatography (right panel). The protection by TpiX suggests an interaction between the toxin TpeX and its cognate immunity protein.

We then modelled the possible interaction between TpeX and its immunity protein, TpiX, using AlphaFold 3. The five predicted TpeX-TpiX complexes (Fig. 3C) displayed a pTM score of 0.70, but a low ipTM score of 0.29, indicating limited confidence in the predicted protein-protein interface. Nevertheless, the five models converged towards a similar overall arrangement, consistently positioning residues 170-180 of TpiX within a predominantly hydrophobic pocket in the N-terminal domain of TpeX (Fig. S3). This recurrent positioning suggests that residues 170-180 of TpiX may contribute to its interaction with TpeX, although the predicted interface remains to be experimentally validated.

All together, these data allow us to conclude that TpiX is the immunity protein of TpeX.

### TpeX dissipates membrane potential of target cells and disrupts their membrane integrity

The membrane potential, a component of the proton-motive force, is an electrical potential across the membrane, and a source of free energy that enables cells to perform chemical and mechanical work^51^. To test our hypothesis that the inner membrane is the target of TpeX and the source of TpeX-mediated toxic activity demonstrated above (Fig. 2B), we used DiOC_2_(3) (3,3-diethyloxacarbo-cyanine iodide) as membrane potential-sensitive fluorescent dye. Being positively charged, DiOC_2_(3) accumulates within cells in a charge-dependent manner. Red fluorescence emission is a marker of normal membrane potential with a shift to green fluorescence emission in case of membrane potential dissipation^52,53^. CCCP (Carbonyl cyanide m-chlorophenyl hydrazone), an uncoupler that dissipates the proton gradient, was used as a positive depolarizing control. Labelled cells were analyzed by flow cytometry with red/green fluorescence ratio monitored as an indicator of membrane potential state^36^. In *E. coli* liquid culture producing TpeX targeted to inner membrane (SS-TpeX), red/green ratios were significantly reduced (p value <0.001) compared with the negative empty vector control strain (Fig. 4A, compare lanes 1 and 3) indicating that SS-TpeX dissipated the membrane potential of cells. In this assay, the amplitude of SS-TpeX effect was not significantly different from that of the uncoupler CCCP (Fig. 4A, compare lanes 3 and 4). The red/green ratio of the cytoplasmic version of TpeX was not significantly different from that of the negative empty vector control indicating that it does not impact the membrane potential (Fig. 4A, compare lanes 1 and 7). Consequently, we can correlate the SS-TpeX-mediated growth inhibition activity observed above with the dissipation of membrane potential induced by SS-TpeX production.

**Fig. 4.**
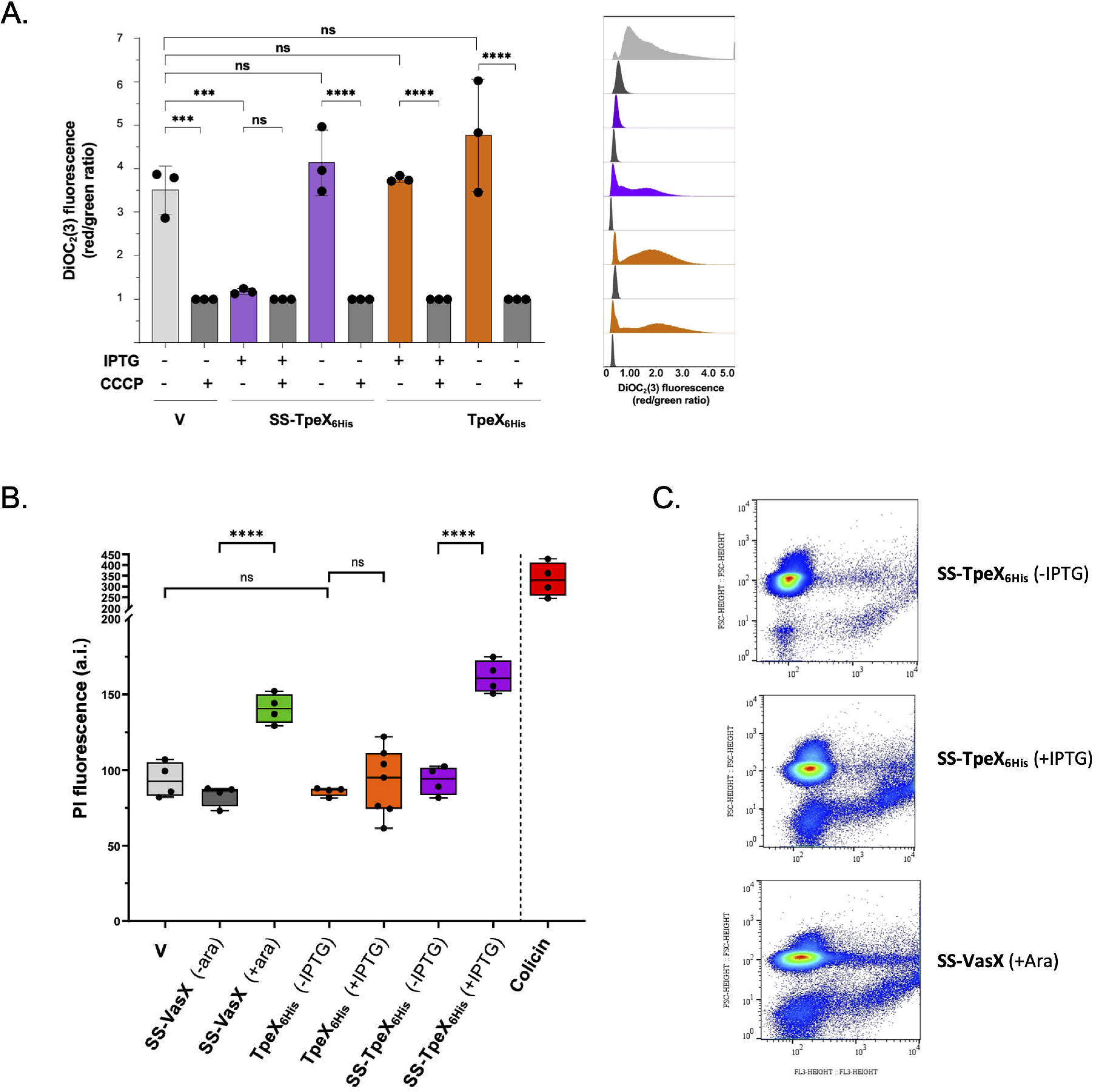
TpeX dissipates membrane potential and impacts membrane integrity. **(A)** *E. coli* BL21(DE3) pLysS producing an inner membrane targeted toxin (SS-TpeX_6H_i_S_ from pSBC106) or a cytoplasmic toxin (TpeX_6His_ from pSBC102) were analyzed with the DiOC_2_(3) probe using flow cytometry. *E. coli* harboring an empty vector (V) was used as negative control. The fluorescent dye DiOC_2_(3) was used as a marker of membrane potential. The red/green fluorescence intensity ratio (615±12.5 nm / 525±15 nm) of DiOC_2_(3) was calculated for each condition. A low red/green ratio characterizes cells in which the membrane potential has been dissipated. The CCCP uncoupler that dissipates the proton gradient was used as a reference and as a positive depolarizing control. Data shown in the left panel represent the mean of triplicate samples. The data shown in the right panel represent the distribution of red/green ratios for cells analyzed in one of the experiment shown on left panel. **(B)** Cells with permeant membranes were analyzed using propidium iodide (PI) staining (22.5 pM). SS-VasX and colicin A (lmg/ml) toxins were used as positive controls for high level of permeant cells (higher PI fluorescence signal) on *E. coli* cells. Cells producing SS-TpeX_6His_ or TpeX_6H_i_S_ are shown in violet and orange respectively. Data shown in the panel B represents the mean of 4 to 7 samples. Statistical analyses were conducted using a two-ways ANOVA Tukey test, ns: non-significant, * p value <0.05, ** p value <0.01, *** p value <0.001, **** p value <0.0001. **(C)** Density plots (Forward scatter FSC versus PI fluorescence) for *E. coli* producing SS-TpeX_6H_i_S_ (from pSBC106, + IPTG) or not (-IPTG), or SS-VasX (from pPER5, + arabinose) showing more cells debris in the presence of an active toxin.

Among T6SS PFTs affecting the membrane potential, VasX (*V. cholerae*), Tse5 (*P. aeruginosa*), Tme1/2 (*Vibrio parahaemolyticus*), TpeV (*V. cholerae* BGT49) form large non-selective pores^24,36,54^ whereas Tse4 (*P. aeruginosa*), Ssp6 (*S. marcescens*), Tke5 (*Pseudomonas putida*) form ion selective ionophores that do not impact membrane integrity^23,44,55^.To investigate the type of pore that could be generated by TpeX we used propidium iodide (PI) labelling. PI is an intercalating agent and fluorescent molecule that binds DNA. Because it is excluded from cells with intact plasma membranes, it is used to evaluate cell viability and integrity. Labelled cells were analyzed by flow cytometry associated with red fluorescence monitoring (Fig. 4B). As PI is membrane impermeant, an increase in red fluorescence is indicative of PI diffusion across a disrupted membrane. We used cells producing SS-VasX^54^ or incubated with colicin A (1mg/ml), two well characterized PFTs, as positive controls for PI diffusion. As expected, a significant increase in PI fluorescence in comparison to negative empty vector control strain was observed with both proteins (Fig. 4B, compare lane 1 with 3 and 8). Whereas production of cytoplasmic TpeX did not significantly increase red fluorescence (Fig. 3B, compare lanes 4 and 5), production of SS-TpeX targeted to inner membrane (Fig. 4B, compare lanes 6 and 7) increased significantly red fluorescence (p value <0.0001) indicating a permeability to PI. Like the two PFTs VasX and colicin A, TpeX affects membrane integrity of bacterial cells most likely by large pores formation.

### TpeX is a bactericidal pore-forming toxin

We took advantage of the heterologous toxicity assay (Fig. 2B) to determine whether TpeX has a bacteriostatic or bactericidal impact on cell growth and viability. The aim of this experiment was to observe whether bacteria can resume growth (bacteriostatic effect) or not (bactericidal effect) after the toxic form of TpeX was produced in *E. coli*. Bacteria were grown in liquid medium harvested 60-minutes post-induction of *ss-tpeX* or *tpeX,* washed and, after normalization to OD_600_, spotted onto a repressive LB agar medium according to the protocol of Bartoli *et al*., 2023^56^. In figure 5A, bacteria harvested before the induction of *ss-tpeX* (t0) were able to grow on the repressive medium, similar to the TpeX or negative control strain. After 60 min of SS-TpeX production, whereas the protein is only weakly detected (Fig. 5A lower panel t60), the bacteria were no longer able to resume growth (Fig. 5A upper panel t60). Taken together, these results indicate that TpeX is a bactericidal toxin.

**Fig. 5.**
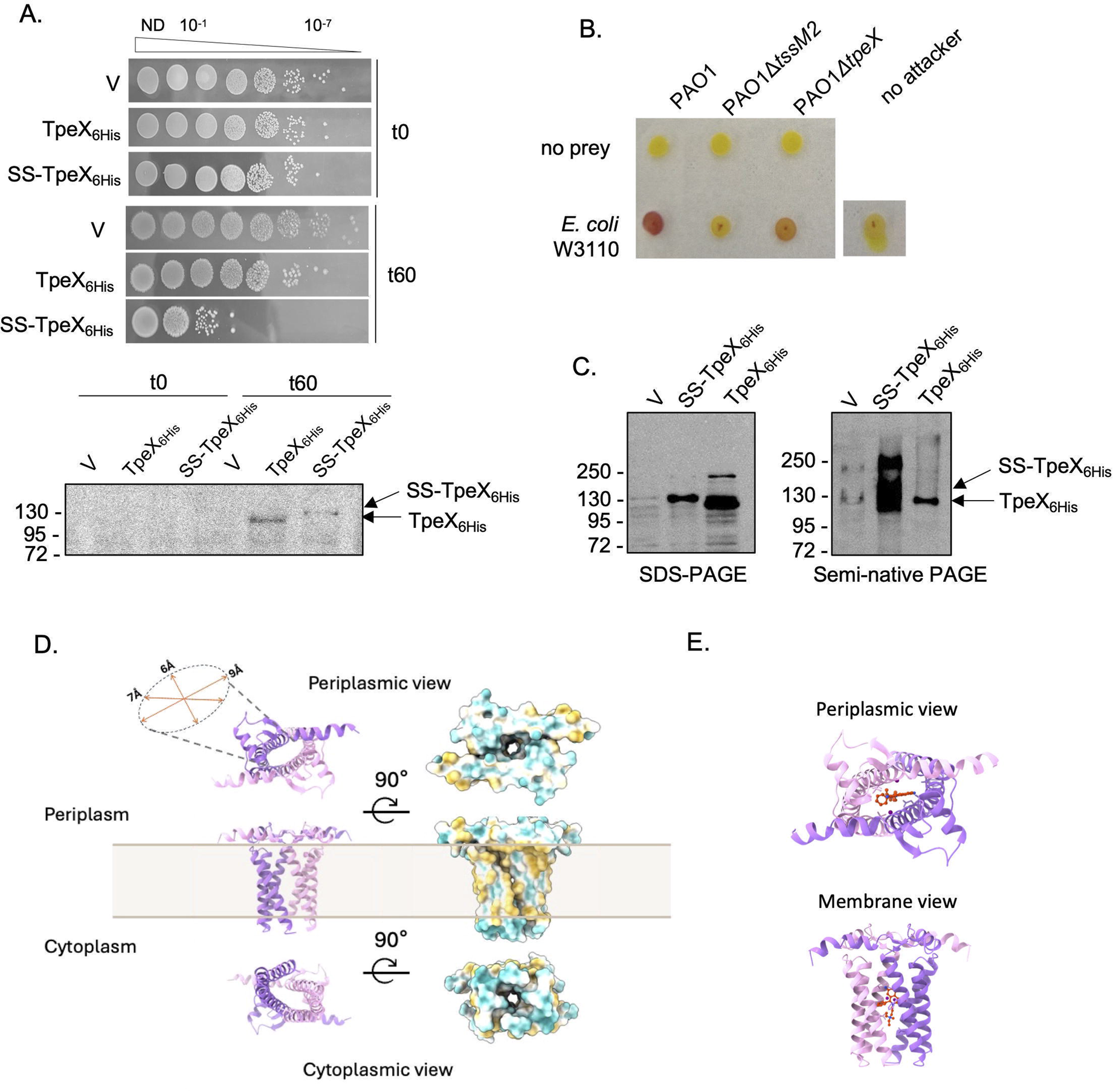
Bactericidal effect through cell lysis upon SS-TpeX oligomerization in the membrane. **(A)** *E. coli* BL31(DE3) pLysS cells from panel (2B) were harvested 0 and 60 min post-induction. Serial dilutions of normalized cultures were spotted on LB agar containing 1% glucose to repress the production of the indicated protein (upper panel). SS-TpeX_6His_ and TpeX_6His_ immunodetection in those samples (bottom panel). **(B)** *E. coli* W3110 was co-cultured for 4 h at 37°C with PAO1 and its isogenic mutants *Etssm2* or *EtpeX* at a ratio of 4 prey for 1 attacker. CPRG at 4 mM was then added to visualize the P-galactosidase activity. The same strains, grown separately (no prey, no attacker), serve as lysis controls. **(C)** *E. coli* BL31(DE3) pLysS cells producing SS-TpeX_6His_ or TpeX_6His_ were analyzed by SDS-PAGE or semi-native PAGE, followed with immunoblotting with anti-TpeX antiserum. The bacterial culture equivalent to 0.2 units of OD_600_ was loaded on a 10% acrylamide gel. The position of the proteins and the molecular mass markers (in kDa) are indicated. **(D)** TpeX dimer prediction by AlphaFold 3. left: presentation by chain (purple for one protomer of TpeX, pink for a second), right: hydrophobicity model of the complex (yellow and blue colours indicate respectively the surface of hydrophobic or hydrophilic residues). **(E)** Prediction of propidium iodide (in red) interaction with the TpeX pore by Boltz-2.

To confirm the role of TpeX in interbacterial competition, we used a colorimetric lysis-associated β-galactosidase assay (LAGA) to monitor the attack by *P. aeruginosa* of *E. coli* prey bacteria^57^. If the antibacterial activity of the attackers results in prey lysis, β-galactosidase is released into the medium. Its extracellular activity is then determined by the hydrolysis of the chlorophenol-red β-D-galactopyranoside (CPRG), which turns from yellow to red. *E. coli* W3110 prey bacteria were co-cultivated with various *P. aeruginosa* attackers, the wild-type (WT) parental strain, the Δ*tssM2* (H2-T6SS mutant) or the Δ*tpeX* isogenic mutants, for 4 hours at 37°C under the H2-T6SS antibacterial conditions we and others previously determined^13,58^. As shown in Figure 5B, the co-culture spot with the WT strain turned red, indicating lysis of the prey. In contrast, the co-culture with the H2-T6SS mutant or the prey alone remained yellow. Co-culturing with the Δ*tpeX* mutant produced an orange color, suggesting that TpeX plays a significant role in lysis together with other effectors. In line with this, we noticed a higher level of cells debris in the presence of SS-TpeX or SS-VasX compared to the negative control (Fig. 4C). Lysis could also result secondarily from membrane potential dissipation^34^. As mentioned earlier, bacterial PFTs undergo structural transformations from soluble and inactive monomers to active and multimeric transmembrane pores^42^. To investigate whether TpeX is capable of oligomerization, we compared its behavior after electrophoresis under semi-native and denaturing conditions. Figure 5C shows that the cytoplasmic form of TpeX is detected at the size of its monomer under both conditions. However, SS-TpeX is capable of multimeric assembly, forming at least dimers under semi-native conditions. These data are consistent with the cellular localization determined in Figure 2A and indicate that artificial targeting of TpeX to the membrane via a signal sequence mimics its insertion into the membrane of target bacteria.

To model the TpeX transmembrane dimer, we performed AlphaFold 3 predictions using the TMH domain (residues 714-838). When the TMH domain was modelled alone, the predicted dimer displayed a very low ipTM score (0.17), indicating low confidence in the predicted interchain interface. Inclusion of hydrophobic or polar ligands markedly increased the confidence of the predicted dimer, with citrate yielding the most consistent models. In the presence of 2 to 6 citrate molecules, the predicted complexes displayed comparable ipTM scores (≈0.82) and high pTM scores (0.84-0.86) (Fig. 5D, left, and Fig. S4). The selected model containing two citrate molecules displays multiple interchain contacts distributed along the transmembrane region, supporting the overall organization of the two TpeX chains within the predicted helix bundle (Fig. S4). Because increasing the number of citrate molecules beyond two provided no substantial improvement in prediction confidence and could introduce an unsupported assumption regarding ligand stoichiometry, we selected the model containing two citrate molecules for further analysis. This model represents a parsimonious solution that provides a high-confidence dimer interface without imposing an unsupported ligand stoichiometry. Importantly, models generated with larger numbers of citrate molecules exhibited highly similar overall architectures, suggesting that the helix-bundle organization primarily reflects intrinsic properties of the TpeX transmembrane domain rather than a specific citrate stoichiometry. The hydrophobicity surface of the selected model (Fig. 5D, right) indicates that the outer surface of the transmembrane bundle is predominantly hydrophobic, whereas the central cavity is more hydrophilic at both the cytoplasmic and periplasmic entrances. The diameter of this central cavity is predicted to range from approximately 6 to 9 Å along the membrane-spanning region. Together, these features define a membrane-spanning cavity within the predicted TpeX dimer, although experimental data will be required to establish whether this predicted cavity corresponds to a functional pore.

As the entry of PI may result not only from the membrane lysis but also from its passage through the pores formed by TpeX, we then modeled the TpeX pore in complex with PI using Boltz-2 (Fig. 5E). The resulting model was highly confident (pTM ≈ 0.91, ipTM ≈ 0.88), with PI consistently positioned within the pore lumen. In addition, the low predicted alignment error (PAE) values at the protein-ligand interface support a specific interaction rather than a nonspecific membrane association. Altogether, these structural models are in good agreement with our biological data (Fig. 4B) and support the idea that PI can be accommodated within the TpeX pore.

In summary, the bactericidal activity of TpeX results in lysis of the prey bacterium and is based on at least the dimerization of TpeX in the cytoplasmic membrane of the latter.

### The complex between TpeX and the HcpB-VgrG6 arrow

The *tpeX* gene is located downstream of the *hcpB* and *vgrG6* genes, which encode components of the T6SS perforating tip, and according literature of the H2-T6SS of PAO1^14,59^. Moreover, the dependence of the Tpx2 pore-forming effector from *P. aeruginosa* PA14 on VgrG6 and H2-T6SS was demonstrated by intra-*Pseudomonas* competition assays and co-purification^32^. To support the delivery of TpeX through the HcpB-VgrG6 arrow we predicted by AlphaFold 3 the structure of the tripartite complex as we did recently for the Tle1 effector^48^ (Fig. 6A). We obtained a confident model of a complex composed of an Hcp hexamer, on which is placed a VgrG6 trimer (each protomer being annotated VgrG6 (1), VgrG6 (2) and VgrG6 (3)), and a TpeX monomer, supported by the Alphabridge score (0.74), piCS (predicted interaction Confidence Score) and pLDDT (predicted Local Distance Difference Test) scores, and PAE (Predicted Aligned Error) plots (Fig. 6B, 6C and 6D). The predicted model is coherent with a T6SS tip composed by a VgrG6 trimer, lying on an HcpB tube made of hexamers and carrying the TpeX effector. Interaction is predicted between the N-terminus of TpeX (amino acids 4-290), and the C-terminus of VgrG6 (amino-acids 667-691) (Fig. 6B & 6D). As said before, the MIX motif of TpeX is present at the N-terminus (amino-acids 1-165, Fig. 1C), but in the model, only one residue, the Glutamine at position 40 of the first MIX motif is predicted to interact with the Leucine 360 of VgrG6 (Fig. 6B). Remarkably, VgrG6 residues 667 to 693 and 682 to 685 form respectively a 7-residue and a 4-residue β-strand upon binding with TpeX that intercalates with a β-sheet of TpeX N-terminus (indicated with black arrows in the left insert of Fig. 6A). The interaction of the N-terminus of TpeX with VgrG6 might thus be stabilized by conformational changes at the C-terminal end of VgrG6. Interestingly, this region of TpeX predicted to interact with VgrG6 overlaps, at least partially, with the surface involved in the TpeX-TpiX complex model (Fig. 3C).

**Fig. 6.**
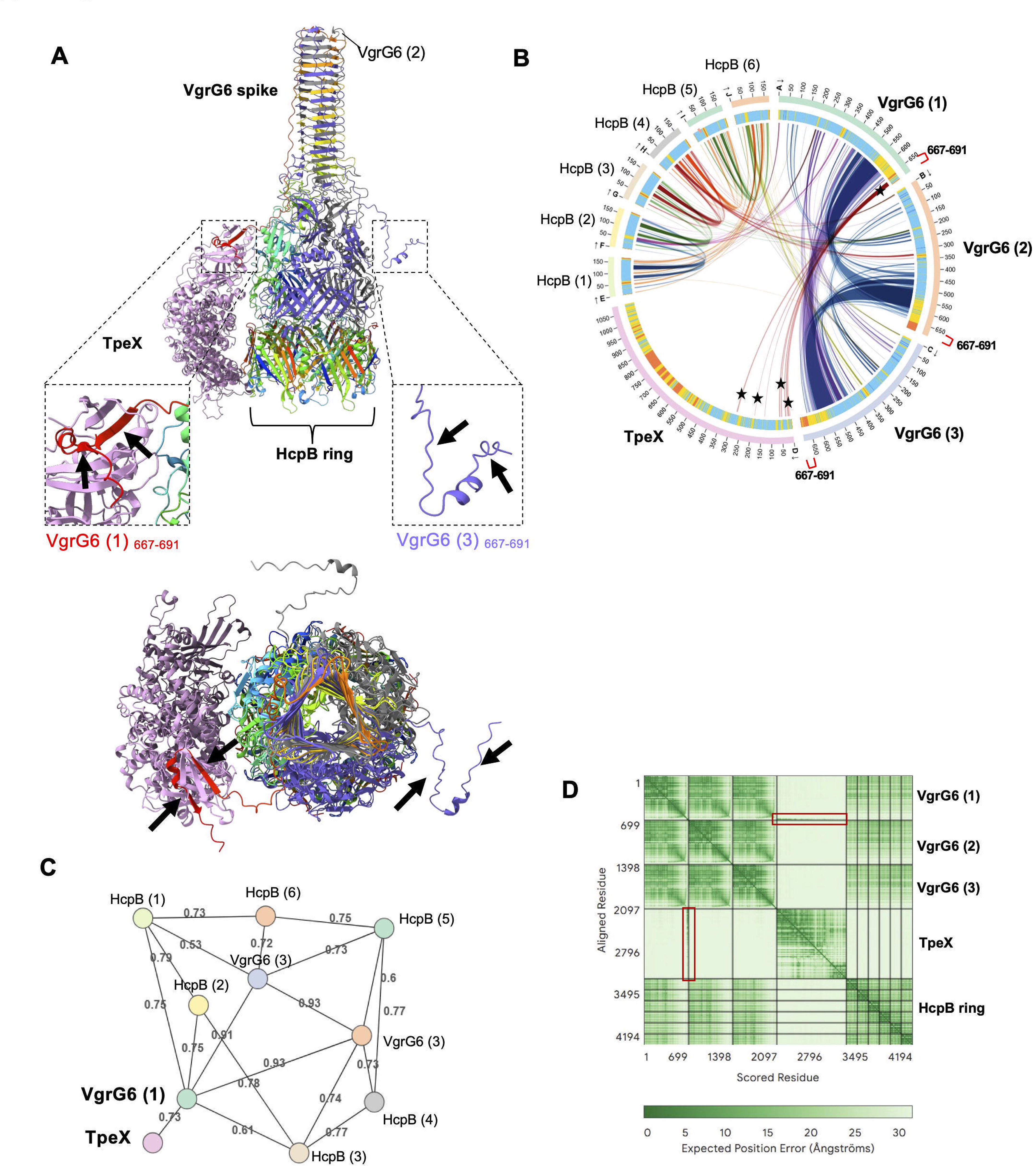
Loading of TpeX on the VgrG6-HcpB spike. **(A)** AlphaFold 3-predicted structure of the complex formed by a VgrG6 trimer, a TpeX monomer and a HcpB hexamer. Top figure: front view, bottom figure: top view. Inserts highlight the structured distal end of VgrG6 interacting with TpeX (protomer 1) (left) and the unstructured distal end of VgrG6 (protomer 3) (right). The black arrow indicates the seven-residue p strand (667-673) and the four-residue P strand (682-685) (insert 1) formed upon interaction with TpeX, which are absent in VgrG6 (insert 3). **(B)** AlphaBridge diagram of the predicted complex. The outer and inner rings represent the number of residues and the pLDDT score of each chain, respectively. The colors of the outer ring correspond to pLDDT confidence levels: blue (very high), cyan (high), yellow (low) and orange (very low). The regions in contact at the VgrG6-TpeX interfaces are indicated by curves marked with black stars. The last twenty residues of each VgrG are marked with red brackets on the diagram. **(C)** The interfaces between the different proteins are shown with the associated scores identified by AlphaBridge. **(D)** PAE (Predicted Aligned Error) graphs predicted by AlphaFold 3. The colored bar corresponds to the predicted position errors (in A). The red rectangles highlight the areas of high confidence (low error values) corresponding to the interfaces between VgrG6 (1) and TpeX.

## DISCUSSION

Our characterization of TpeX (PA5265) from *P. aeruginosa* PAO1 identifies a VasX-like (Fig. 1), membrane-targeting T6SS effector that dissipates the membrane potential and compromises membrane integrity of prey bacteria (Fig. 4), presumably via oligomerization in the membrane (Fig. 2 & Fig. 5). The bactericidal activity of TpeX that results in cell lysis (Fig. 4 & Fig. 5) can be antagonized by interaction with the cognate immunity protein TpiX (Fig. 3). TpeX differs markedly from previously described *P. aeruginosa* PFTs such as Tse4 and Tse5, two H1-T6SS-dependent effectors of the PAO1 strain. Tse4 is a small protein that creates narrow ion-selective channels coupling cell depolarization with K^+^ efflux^60^ and Tse5 is a <u>r</u>earrangement <u>h</u>ot <u>s</u>pot (Rhs) protein, with an auto-proteolytic product forming large pores^24^. These features position TpeX within the growing class of pore-forming T6SS effectors that operate at the bacterial membrane to mediate interbacterial antagonism.

A recent genomic analysis of the diversity of T6SS effectors in the 1,960 published genomes identified TpeX as an accessory effector of *P. aeruginosa* PAO1, presumably linked to H2-T6SS^16^. In contrast to the H1-T6SS, which is predominantly linked to core effectors, the set of effectors associated with H2-T6SS exhibits high variability in terms of their number (presence or absence) across different loci. This indicates that, depending on the various niches of *P. aeruginosa* strains, the repertoire of effectors, and more particularly of the H2-T6SS machinery, has adapted and diversified. Accessory effectors can be considered as a selective advantage in local ecosystems or against niche-specific competitors. The *tpeX* gene is associated with the orphan *vgrG6* locus (Fig. 1). A unique feature of this locus among *P. aeruginosa* strains is the presence of mutually exclusive effectors, TpeX (TspE1b) and Ptx2 (TpsE1c)^16^. Like TpeX, Ptx2 is a VasX-like effector of the PA14 strain. Recent work on Ptx2^32,33^ has shown that it is an H2-T6SS dependent effector and requiring both an immunity protein Pti2 and a cytoplasmic DUF4123 family adaptor Tap6, like VasX yet unlike TpeX. The interaction of Ptx2 with the C-terminus extremity of VgrG6 through the adaptor protein Tap6 was studied by co-purification and structural model prediction.

Our biochemical and cell-based data complement the observations regarding Ptx2. Indeed, the ability of SS-TpeX to assemble into multimers (Fig. 5) and to depolarize membranes (Fig. 4) mirrors behaviors described for VasX and other pore-forming effectors and supports a model in which oligomerization at the target membrane is a key step in pore formation and bactericidal activity. That truncated colicin-like fragments of TpeX retain partial toxicity (Fig. 2B) suggests that the C-terminal domain is the minimal membrane-disrupting unit, whereas the N-terminal domain likely mediates targeting to the membrane, as shown for VasX^36^. The substitution of two residues in the transmembrane helices of Ptx2 effectively demonstrated that this region is responsible for toxicity, since those variants were not toxic anymore^33^. As demonstrated for Ptx2, TpeX should be dependent on H2-T6SS of PAO1. Indeed, the orphan *vgrG6* locus of PAO1 has been linked to the H2-T6SS by a phylogenetic clustering and co-regulation^14^. Furthermore, VgrG6 is required for the delivery of TseT, a known H2-T6SS effector in PAO1, which gene is located at a distance from *vgrG6*^59^. Our LAGA assay, which demonstrates TpeX-dependent lysis of prey bacteria, was performed under conditions where the H2-T6SS machinery is active, since a *tssM2* mutant no longer lyses *E. coli* (Fig. 5B). This provides indirect evidence supporting a link between TpeX and the H2-T6SS machinery. To further support the delivery of TpeX through the HcpB-VgrG6 arrow we predicted by AlphFold 3 the structure of the tripartite complex (Fig. 6A). The model proposes a T6SS tip composed by a HcpB hexamer supporting a VgrG6 trimer carrying the TpeX effector. In contact with TpeX, two β-strands are predicted to form at the C-terminus of VgrG6 and to intercalate with a β-sheet of the MIX motif at the N-terminus of TpeX. At the contrary, VasX and Ptx2 need an adaptor of the DUF4123 family, called VasW and Tap6 respectively^32,34^. As shown by Habich *et al.* (2025)^16^, the absence of an adaptor is a trait of PAO1-like strains containing *tpsEb* (*tpeX*) effector gene. This is a sign of a structural evolution of TpeX and selection of a structure that allows secretion without an adaptor. In PA14, the C-terminus of VgrG6 recruits Ptx2 via a helix-turn-helix motif, while the Tap6 adaptor interacts with both, VgrG6 and Ptx2^32^.

AlphaFold 3 predicts the same for VasX and its VasW adaptor protein (Fig. 1 and Fig. S5).Thus, Ptx2 and VasX would require Tap6 and VasW to establish a stable interaction with their cognate VgrG, as we recently showed with Tle1 and its adaptor protein Tla1 with VgrG4a^48^. In contrast, TpeX does not appear to require an adaptor, as the intercalation of the 2 β-strands of VgrG6 into the β-sheet of TpeX probably ensures the stabilization of the complex on its own. These convergent observations across PAO1 and PA14 argue for a two-tiered strategy: precise effector delivery via adaptor-VgrG recognition, and robust local neutralization by membrane-associated immunity to avoid sister cells attack.

Even if pore architectures are variable among PFTs, two large groups are considered, α-PFTs and β-PFTs, assembling to membrane from secondary structure based α-helices or β-barrels domains respectively ^42^. Regarding our results, TpeX belongs to α-PFTs from the colicin family. All described PFTs transition from a soluble inactive monomer to a toxic transmembrane oligomeric assembly forming a pore^42,61^, from one topological fold to another one. Regarding TpeX, the first interaction could be mediated by the TpeX N-terminal predicted PH domain with membrane lipids (Fig. 1D), increasing TpeX local concentration and favorizing oligomerization required for pore formation in the lipid amphipathic bilayer^42,62^. Following insertion, hydrophilic residues are exposed in the lumen of the pore region, whereas hydrophobic regions are exposed towards the fatty acid tails in the bilayer^61^ (Fig. 5D). According to our results, TpiX could bind directly TpeX to prevent pore formation and toxicity from sister cells (Fig. 3). In line with this, to study intra-species competition, Rudzite *et al.* (2023)^63^ were able to delete the *tpiX* gene by also deleting *tpeX*. Since the genes encoding immunity are essential genes, as previously observed for *tpiX*^45^, the viability of this mutant supports the role for TpiX as an immunity protein of TpeX. Remarkably, our predictions concerning the formation of complex structures, between TpeX and TpiX, and between TpeX and VgrG6 and HcpB, would involve the N-terminal region of TpeX, from residues 56 to 197 for TpiX (Sup. Fig. S3) and from 4 to 290 for VgrG6 (Fig. 6). These structures would include the MIX motif (residues 1-165, see Fig. 1C), but only one residue from the first MIX motif could interact with a residue from VgrG6, none from TpiX (Gln_40_ of TpeX with Leu_360_ of VgrG6; Fig. 6B & Fig. S3). These predictions do not allow us to draw any conclusions regarding the importance of the MIX motif for VgrG targeting. Based on our various observations, we propose a mechanistic hypothesis whereby TpiX could stabilize TpeX in a soluble, inactive conformation by occupying the N-terminal region, thereby preventing the structural rearrangement required for its insertion into the membrane. In this model, VgrG could also help to maintain TpeX in a soluble form prior to its delivery, while its dissociation would subsequently allow TpeX to adopt its active conformation in the membrane. However, this model remains hypothetical, particularly given the low confidence in the TpeX-TpiX interface predicted by AlphaFold 3. The convergence of the five models and the observed overlap with the VgrG interaction region therefore constitute evidence in favor of a mechanistic model to be tested experimentally, rather than a direct demonstration.

Taken together, our data position TpeX as a potent, oligomerization-dependent, membrane-disrupting T6SS effector of PAO1 that uses a classical toxin-immunity pair to mediate interbacterial antagonism. Because the membrane potential governs a wide range of bacterial physiology and behaviors, for example ATP synthesis, pH homeostasis, metabolism, membrane transport global energy metabolism, cell mobility or cell division^51^, inducing membrane depolarization represents an effective strategy to compete with other microorganisms, and could be extended to eukaryotic membranes. Pore formation is an offensive weapon or a defense strategy conserved through a structural homology among bacteria and eukaryote kingdoms. Indeed, the PH domain of TpeX (Fig. 1D) resembles that of eukaryotic proteins. These PH domains, which are very common in eukaryotic proteomes, were initially discovered to enable interaction with membranes via an interaction with phosphoinositides, and are now also recognized for other functions, notably protein-protein interactions^43^. In *V. cholerae*, VasX has been shown to be a trans-kingdom toxin that impacts both other bacteria and the eukaryotic host. Similar toxins have been described in *P. aeruginosa*^11,13^, raising the question of whether this is also the case for TpeX. The precise molecular mechanisms underlying TpeX pore formation and/or membrane disruption remain to be determined. To fully understand TpeX mode of action, precise oligomeric stoichiometry, architecture, and lumen size need to be characterized. TpeX thus constitutes a highly specific potential therapeutic target for an anti-virulence strategy aimed at combating *P. aeruginosa* infections.

## MATERIALS AND METHODS

### Bioinformatic analysis

Bioinformatics DNA sequences were retrieved from the *Pseudomonas* Genome Database^64^. DNA and amino acid sequence searches were performed using NCBI CDD (Conserved Domains Database)^65^, BLAST (Basic Local Alignment Search Tool)^66^ and Phyre2 (PHYRE2 Protein Fold Recognition Server)^67^. Secondary protein structure, topology, subcellular localization and signal peptide were predicted using transmembrane helices prediction TMHMM (TransMembrane Hidden Markov Model) V2.0^65,66,67,6868,6969,7068,69^, HHpred server^68^, PSORTb V3.0^70^ and SignalP V6.0^71^. Structural predictions for TpeX and VasX (uniprot codes Q9HTT4 and Q9KNE5 respectively) were obtained from the AlphaFold 2 database. Based on these predictions, we searched for the existence of structural homologs using the Foldseek search server^72–74^ . The analysis was then carried out using UCSF ChimeraX. (v1.8) software. To predict the 3D structures of protein complexes, the amino acid sequences of the proteins of interest, obtained in FASTA format from *Pseudomonas.com*, were submitted to AlphaFold3 (https://alphafoldserver.com/welcome)^75^. These predictions were produced with predicted aligned errors (PAE) and pLDDT values as confidence scores. The AlphaFold-predicted structures were subsequently analyzed with AlphaBridge^76^, investigate residue-residue interactions at the predicted interfaces. Confidence in these predicted interfaces was assessed using the predicted interaction confidence score (piCS), with a default cut-off of 0,7. Boltz-2 (v2.2.0)^77^ via Tamarind Bio was used to predict the structure of protein-ligand. The ligand was defined based on its SMILES representation from PubChem (CC[N+](C)(CC)CCC[N+]1=C2C=C(C=CC2=C3C=CC(=CC3=C1C4=CC=CC=C4)N)N.[I-].[I-]).

The AlphaFold 3 structure of the dimer was used as a template to preserve the quaternary architecture. Predictions were made with 5 recycling cycles and physical potentials enabled. Fifteen independent models were generated, and the one with the highest confidence score was selected. The structures were analyzed using UCSF ChimeraX (v1.8).

### Bacterial strains, plasmids and growth conditions

All strains of *E. coli* and *P. aeruginosa* and used in this study are listed in Table 1. The E*. coli strains* CC118λPir and K-12 DH5α were used for cloning procedures, while BL21(DE3) pLysS was employed for protein expression/production under T7 promoter. Cultures were grown in LB medium at 37°C, with specific growth conditions provided in the main text when required. Plasmids were introduced into *P. aeruginosa* through triparental mating, facilitated by the helper plasmid pRK2013 (Table 1). Plasmid maintenance was ensured by supplementing media with appropriate antibiotics: ampicillin (50 μg/ml) for *E. coli*, kanamycin (50 μg/ml) for *E. coli*, streptomycin (30 μg/ml for *E. coli* and 2,000 μg/ml for *P. aeruginosa*).

**Table 1:**
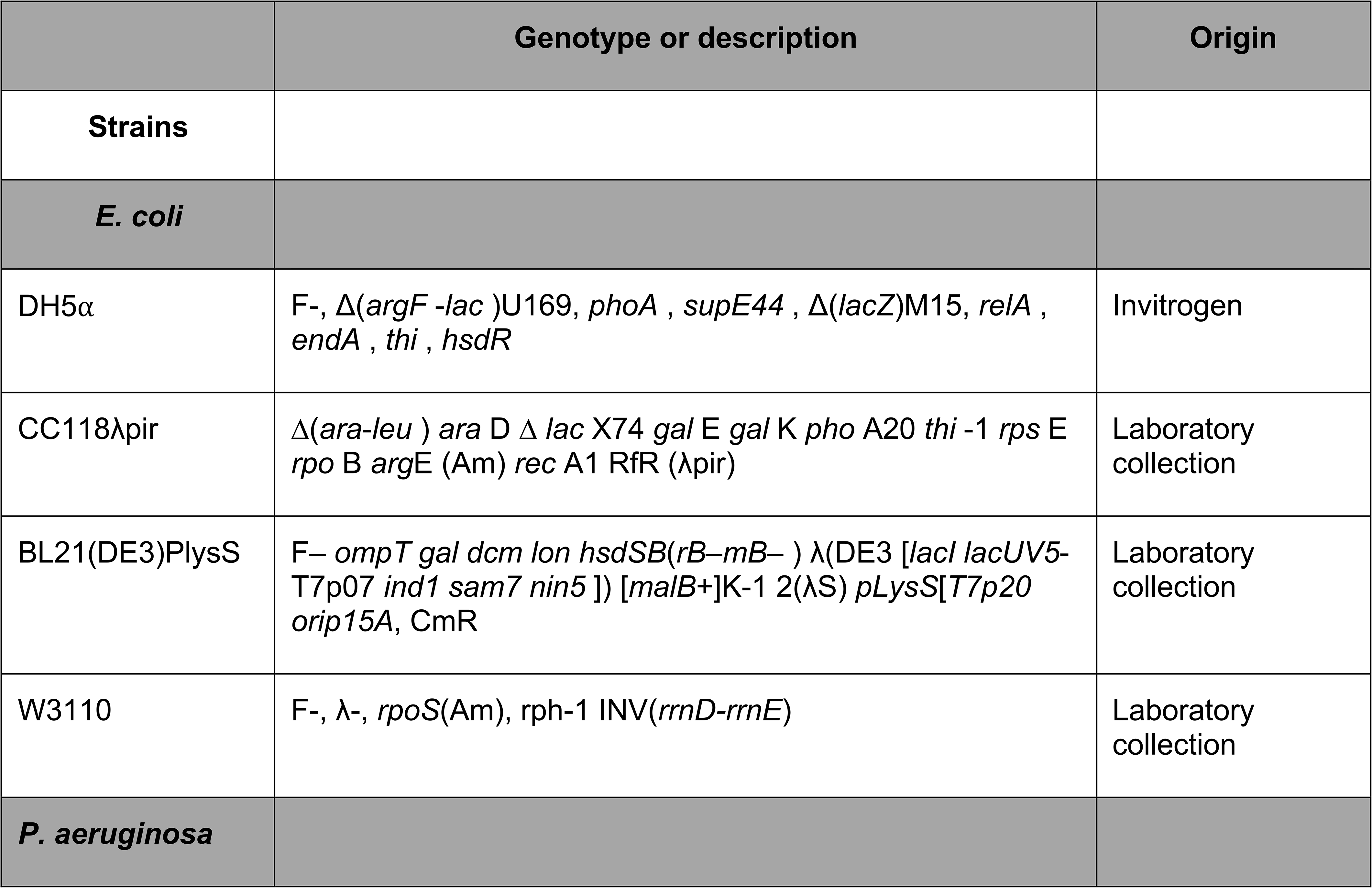
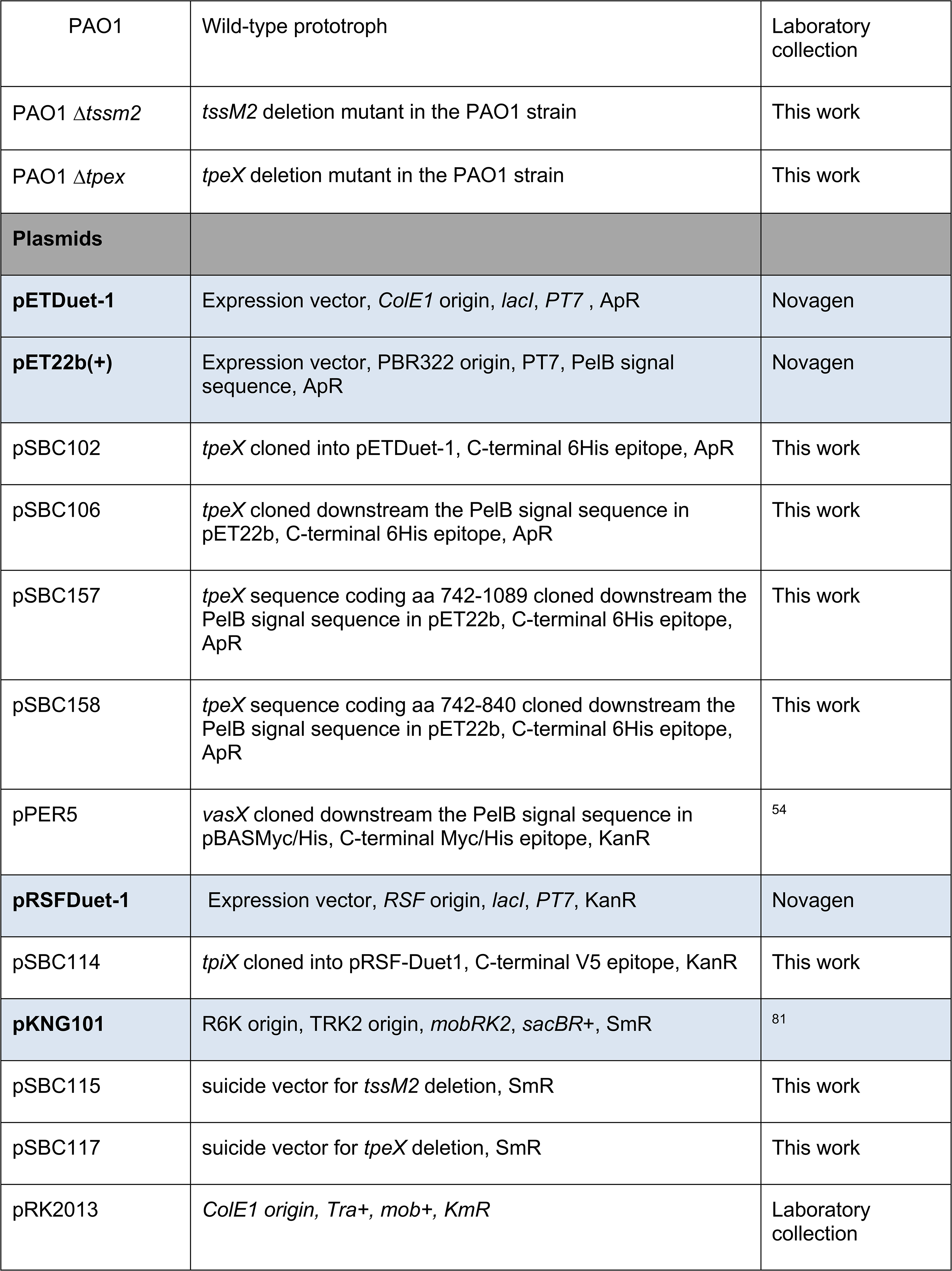
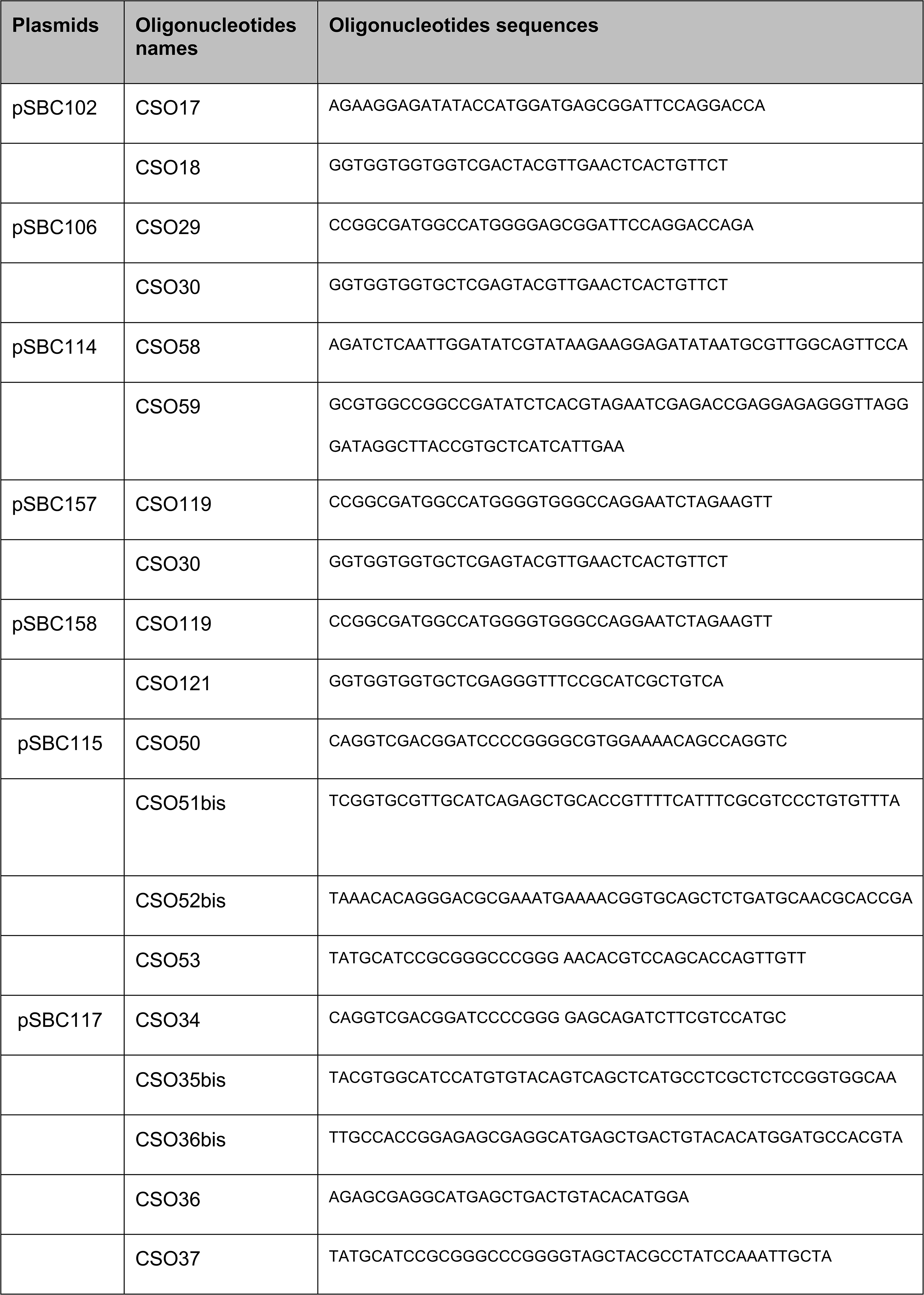
*E. coli* and *P. aeruginosa* strains and plasmids used in this study.

Cloning was performed by sequence and ligation independent cloning (SLIC^78^) and cloned sequences were confirmed by DNA sequencing. Table 1 provides a list of plasmids constructed and used, and the oligonucleotides synthesized by Eurogentec and IDT. During the selection steps of clones potentially carrying a recombinant plasmid encoding a toxic form of TpeX, glucose (0.4%) was systematically added. The construction of the Δ*tssM2* and Δ*tpeX* mutants was performed as described in Berni *et al*., 2019. The PCR fragments corresponding to 500 bp upstream and 500 bp downstream *tssM2* and *tpeX* and designed to avoid any polar effect, were cloned into the pKNG101 suicide vector. After sequence verification, the resulting pKNG101 constructs were transferred into *P. aeruginosa* via conjugation. The mutants, in which the double recombination events occurred, were confirmed by PCR analysis.

### Subcellular fractionation

Fractionation of *E. coli* cells was done as described previously^79^.

### Immunodetection

Protein samples corresponding to equivalent culture densities (measured by optical density at 600 nm) were resuspended in Laemmli loading buffer, boiled, and subjected to SDS-PAGE. The semi-native electrophoresis was performed as described in Salacha *et al.,* 2010^80^. The samples were resuspended in native Laemmli buffer (containing SDS 0,2% and without β-mercaptoethanol) and were not boiled before electrophoresis. The semi-native gels were prepared without SDS, and the electrophoresis buffer was the same as for SDS-PAGE.

Proteins were subsequently detected via immunoblotting as described by Sana *et al.* (2015), using primary monoclonal antibodies against His6 (Penta His, Qiagen, 1:1,000), V5 (Bethyl Laboratories, 1:1,000), TolA (laboratory collection, 1:500), TolB (laboratory collection, 1:500), TpeX (anti-peptides developed by GenScript (this study), 1:500). Peroxidase-conjugated anti-Rabbit IgGs (Sigma, dilution 1:5000) were employed as secondary antibodies. Protein revelation was carried out using a homemade enhanced chemiluminescence and membrane were analyzed with ImageQuant LAS4000 software (GE Healthcare Life Sciences).

### Heterologous toxicity assays

*E. coli* BL21(DE3) pLysS containing plasmids producing targeted proteins were grown overnight at 37°C in LB with 0.4% of glucose. Serially diluted bacterial suspensions (10 μl) were spotted onto LB agar plates containing either 0.1 mM IPTG or 0.4% glucose and incubated at 37°C for 16 hours as described in Berni *et al*., 2019^13^.

### Protein purification by affinity chromatography

*E. coli* BL21(DE3) pLysS cells harboring pSBC102 and pSBC114 were cultivated in LB medium at 37°C until an OD_600_ of 0.5 was reached. The expression of the *tpeX* and *tpiX* genes was induced by adding 0.5 mM IPTG, followed by incubation for 1 hour at 37°C. Cells were then collected by centrifugation at 4,500 rpm for 15 minutes at 4°C and the resulting cell pellets were stored at -20°C. The pellets were resuspended in a lysis buffer containing 40 mM Tris–HCl (pH 8.0), 100 mM NaCl, 10 mM imidazole, 5% glycerol, 0.1% Triton X-100 (Sigma), 1 mg/ml lysozyme, 20 μg/ml DNase I (SIGMA), 20 mM MgCl_2_, and 1 mM phenylmethylsulfonyl fluoride (PMSF). Cells were incubated in this lysis buffer for 1 hour at 4°C with gentle rotation, and after that lysed by sonication. The cell lysates were clarified by centrifugation at 15,000 rpm for 30 minutes. The clarified supernatant was incubated for 1.5 hours with Cobalt resin (Thermo Scientific) pre-equilibrated in binding buffer (40 mM Tris–HCl pH 8.0, 100 mM NaCl, 10 mM imidazole, 5% glycerol, 0,1% Triton X-100). The resin was washed with the binding buffer. Target proteins were eluted using the binding buffer with 50 mM imidazole.

### Membrane potential and integrity assays followed by flow cytometry

*E. coli* BL21(DE3) pLysS harboring empty vector, pSBC106 or pSBC102 for expressing respectively membrane or cytoplasmic forms of TpeX were grown overnight in LB supplemented with antibiotics and glucose (0.4%). Overnight cultures were diluted in selective LB and grown at 37°C with agitation (170 rpm). IPTG induction was performed at OD_600_ = 0.5 and growth maintained one additional hour in the same conditions. Cells were diluted in PBS buffer + glucose. Samples were first analyzed for their light scattering (forward scattering angle FSC *versus* side angle scattering SSC signal): the density plot obtained was first gated on the population of cells and then filtered to remove multiple events. DiOC2(3) (3,3-diethyloxacarbocyanine iodide; final concentration 25 µM) labelling was measured using the ratio of red/green fluorescence (FL3: 615/25 nm / FL1: 525/30 nm) and were quantified in each sample in absence or presence of the uncoupler CCCP (Carbonyl cyanide m-chlorophenyl hydrazone; final concentration 20 µM) as reference for positive depolarizing control. Membrane integrity was analyzed using red PI (propidium iodide 22.5µM) fluorescence (FL3: 615/25 nm) on *E. coli* BL21(DE3) pLysS strains producing TpeX (+/-SS), and SS-VasX from overnight LB solid culture at 37°C. A total number of 300 000 particles were collected per sample and data analyses were carried out with 3-7 biological samples with strains expressing different constructions of TpeX. Data were acquired with a S3e cells sorter (Bio-Rad) using both 488 and 561 nm lasers and were analysed using FlowJo v10.6. Statistical analyses were conducted with Prism.v8.2 to estimate the level of significance using a two-way ANOVA and a Tukey correction. ns: non-significant, * p value <0.05, ** p value <0.01, *** p value <0.001, **** p value <0.0001.

### Bacteriostatic or bactericidal effect of the toxin

This assay has been done as in Bartoli *et al.,* 2023^56^. From stationary-phase overnight cultures, fresh LB medium containing 1% glucose was inoculated to an OD_600_ of 0.01. Bacteria were cultivated at 37°C to OD_600_ = 0.3 in LB medium containing 1% glucose, washed twice with LB before induction of PT7 with 0.1 mM IPTG. At time 0 and 60-min post-induction, an aliquot was recovered and chilled in ice water for 2 min. Cells were pelleted at 6,000 g at 4 °C and re-suspended in ice-cold fresh LB. After normalization to an OD_600_ of 0.5, serial dilutions were done in sterile PBS and spotted on LB agar plates containing appropriate antibiotics and 1% glucose.

### Interbacterial competition by lysis-associated β-galactosidase assay (LAGA)

The assay was performed as described in Taillefer *et al*., 2023^57^. *P. aeruginosa* attacker strains were grown overnight at 25°C in LB medium. The day of the competition, the *E.coli* prey strain W3110 (reporter of LacZ activity) was grown to mid-exponential phase at 37°C in LB medium containing 0,1 mM IPTG to induce *lacZ* gene expression. Attacker and prey cells were then mixed respectively at a ratio 1:4 in 10 μl (corresponding to 0,5 OD_600_) and spotted on LB medium agar plate for 4 hours at 37°C. Negative controls were done by spotting on plate separately attacker and prey strains. 10 µl of 4 mM CPRG (chlorophenol-red β-D-galactopyranoside, Roche) was added directly on bacterial spots. If the antibacterial activity of the attacker results in reporter cell lysis and thus release of β-galactosidase, hydrolysis of the yellow colored CPRG into the purple product chorophenol red (CPR) occurred in few minutes. In absence of lysis, the spot stayed yellow. The intensity of the coloration is correlated with CPR concentration and thus directly proportional to the number of lysed reporter cells and to the attacker antibacterial activity.

## Acknowledgments

We thank members of Bleves-Ize team and members of the LISM, in particular D. Duché and J. Desjardin, for helpful discussions and support. We are grateful to Milena Ghazaryan and Elisa Montredon for their experimental contributions during their summer internship and to M. Ba, I. Bringer, A. Brun and M. Guilbert for technical assistance. We thank Pr D. Salomon for sharing the pPER5 plasmid. We acknowledge UCSF ChimeraX for molecular graphics that is developed by the Resource for Biocomputing, Visualization, and Informatics at the University of California, San Francisco, with support from National Institutes of Health R01-GM129325 and the Office of Cyber Infrastructure and Computational Biology, National Institute of Allergy and Infectious Diseases. The English language was improved with the assistance of AI (DeepL).

## Author contributions

CS, SR, DL, MR, BI, GB and SB designed and conceived the experiments. CS, SR, DL, MR, LS and GB performed the experiments. SB supervised the execution of the experiments. CS, SR, DL, MR, LS, BI, GB and SB analyzed and discussed the data. SR and SB wrote the paper with review/editing from all the authors.

## Funding and additional information

This work was supported by the Aix-Marseille Université (amU), the Centre National de la Recherche Scientifique (CNRS), and a grant from the Agence Nationale de la Recherche (ANR-21-CE11-0028). DL and MR were supported with a PhD fellowship from the French Research Ministry, and DL by the Fondation pour la Recherche Médicale, « grant number FDT202404018453 » for a 4^th^ year.

