## Supplementary material for "The Type VI secretion effector TpeX expands the pore-forming virulence arsenal of *Pseudomonas aeruginosa*": Sup Figures

### Supplementray Figure 1

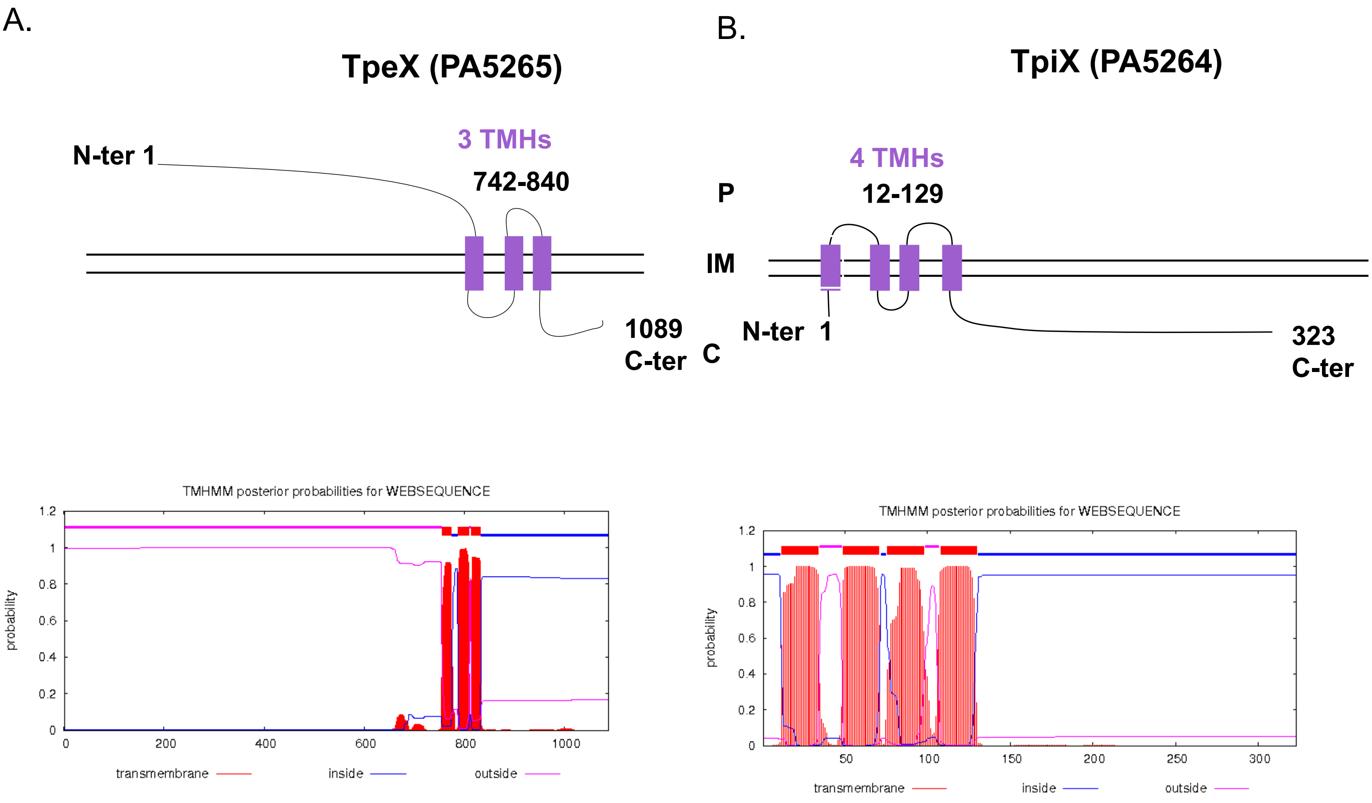


**Fig. S1. PA5265 and PA5264 inner membrane topology prediction. (A)** Schematic representation of PA5265 hypothetical protein (1089 amino acids) predicted to be localized in the inner membrane (IM, PSORTb V3.0) with a large N-terminal (N-ter) domain in the periplasm (P, 1 to 741 ) followed by 3 predicted hydrophobic transmembrane helices (TMHs, 742 to 840) and a C-terminal (C-ter) domain in the cytoplasm (DAS and TMHMM V2.0). **(B)** Schematic representation of PA5264 hypothetical protein (323 amino acids) predicted to be localized in the IM (PSORTb V3.0) with a large N-ter domain in the periplasm (1 to 741) followed by 4 predicted hydrophobic TMHs (AA 12 to 129) and a C-ter domain in the cytoplasm (DAS and TMHMM V2.0).

### Supplementray Figure 2

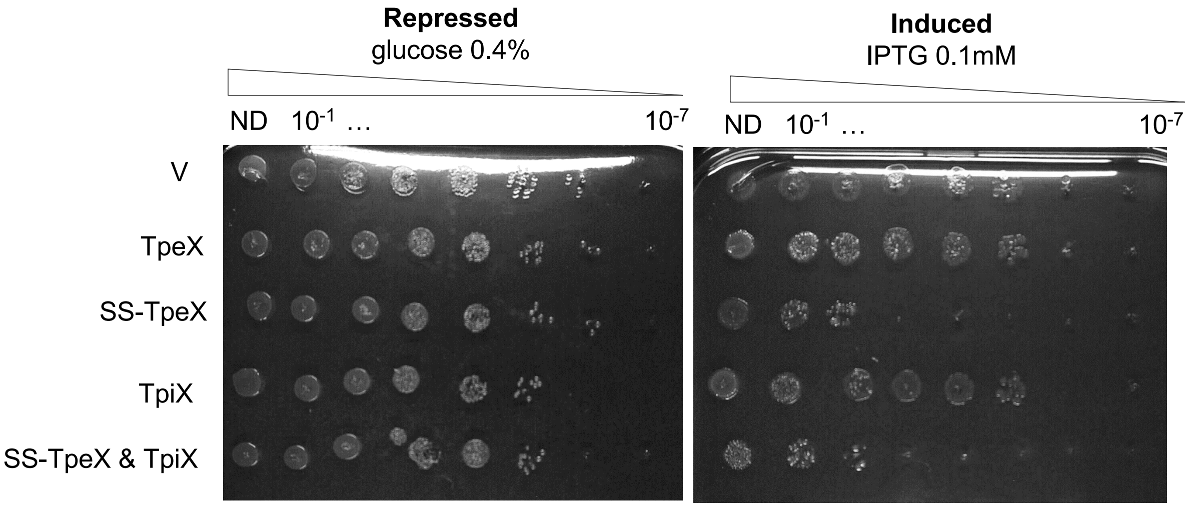


**Fig. S2**. **TpiX does not protect from TpeX toxicity in *E. coli.*** Serial dilutions (from non-diluted to $10^{-7}$) of normalized cultures of *E. coli* BL21(DE3) pLysS producing TpeX (from pSB102) or targeted to the membrane SS-TpeX (from pSB106), or TpiX (from pSBC114) were spotted on LB agar plates supplemented (left panel) with 0.4% glucose or (right panel) with 0.1 mM IPTG. Glucose and IPTG allow respectively repression and induction of the gene encoding the T7 RNA polymerase.

**Supplementray Figure 3**

| **TpeX** | **Distance (Å)** | **TpiX** |
|---|---|---|
| His197 | 4.24 | Leu170 |
| Ala194 | 3.81 | Leu170 |
| Asn190 | 4.08 | Pro172 |
| Val193 | 3.85 | Pro172 |
| Val193 | 4.48 | Ser173 |
| Trp189 | **3.14*** | Ser173 |
| Phe149 | 3.65 | Gly174 |
| Trp189 | **3.42*** | Gly174 |
| Trp189 | 4.15 | Tyr175 |
| Leu144 | 4.15 | Tyr175 |
| Ile56 | 4.33 | Tyr175 |
| Gly147 | 3.74 | Tyr175 |
| Trp189 | 3.54 | Gly176 |
| Val193 | 3.97 | Gly176 |
| Val193 | 4.12 | His178 |
| His197 | 4.45 | Tyr180 |

**Fig. S3**. **Intermolecular contacts between TpeX and TpiX.** Contacts were identified using NCONT with a distance cutoff of 4.5 Å. For residue pairs displaying multiple atom-atom contacts, the shortest distance is reported. Asterisks indicate hydrogen bonds identified by PDBePISA.

### Supplementray Figure 4

**A.**

| AF3 Results | | | |
|---|---|---|---|
| TpeX 714-838 | | | |
| name | number | ipTM | pTM |
| OLA | 50 | 0,6 | 0,63 |
| | 40 | 0,64 | 0,68 |
| | 30 | 0,7 | 0,73 |
| | 20 | 0,66 | 0,71 |
| | 10 | 0,78 | 0,82 |
| | 8 | 0,67 | 0,74 |
| | 6 | 0,71 | 0,77 |
| | 4 | 0,73 | 0,79 |
| | 2 | 0,66 | 0,73 |
| MYR | 20 | 0,7 | 0,75 |
| | 10 | 0,71 | 0,76 |
| | 8 | 0,72 | 0,78 |
| | 6 | 0,74 | 0,8 |
| | 4 | 0,75 | 0,8 |
| | 2 | 0,73 | 0,79 |
| CIT | 10 | 0,77 | 0,82 |
| | 9 | 0,78 | 0,82 |
| | 8 | 0,78 | 0,83 |
| | 7 | 0,81 | 0,85 |
| | 6 | 0,82 | 0,86 |
| | 5 | 0,8 | 0,84 |
| | 4 | 0,81 | 0,84 |
| | 3 | 0,81 | 0,84 |
| | 2 | 0,82 | 0,84 |
| | 1 | 0,77 | 0,8 |
| nothing | 0 | 0,17 | 0,38 |

**B.**

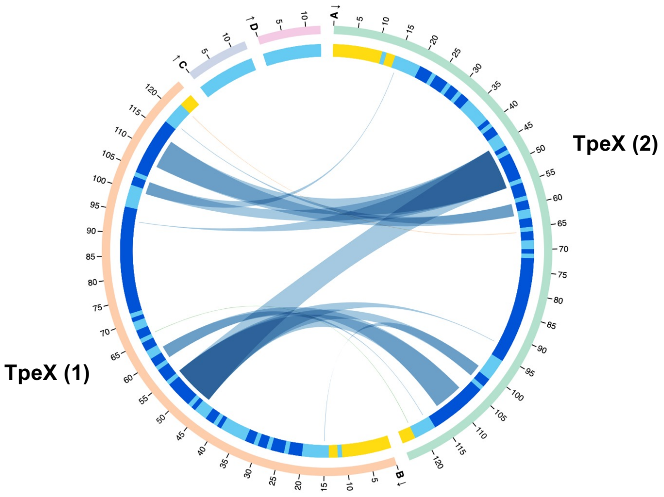


**Fig. S4**. **Modelization by AlphaFold 3 of the TpeX dimer. (A)** AphaFold3 results according ligands utilized to model TpeX dimer. OLA: Oleic acid, MYR: Myristic acid, CIT: citrate. **(B)** The interfaces between the two TpeX protomers are shown with the associated scores identified by AlphaBridge.

### Supplementray Figure 5

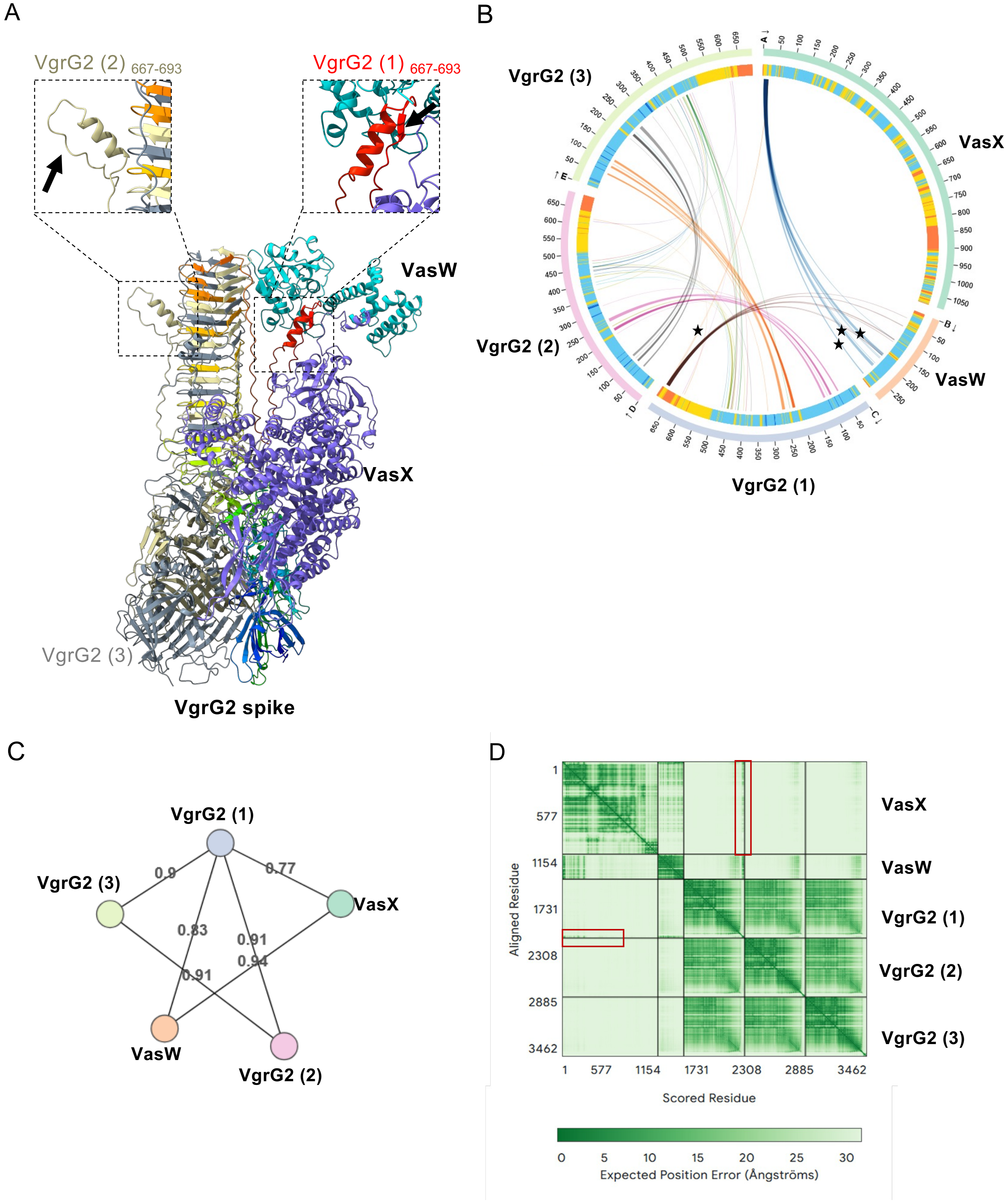


**Figure S5 : The C-terminal domain of VgrG2 recruits VasX and VasW. (A)** Structure predicted by AlphaFold 3 of the complex formed by a VgrG2 trimer, a VasX monomer and a VasW monomer (top: front view; bottom: top view). The insets highlight the structured distal end of VgrG2 (protomer 1) (interacting with VasX) (right) and the unstructured distal end of VgrG2 (protomer 2) (right). The black arrow indicates the β-strand that is absent in VgrG2 (2) and (3). **(B)** AlphaBridge diagram of the predicted complex. The outer and inner rings represent the number of residues and the pLDDT score for each chain, respectively. The colours of the outer ring correspond to the pLDDT confidence levels: blue (very high), cyan (high), yellow (low) and orange (very low). The regions of contact between VgrG2 and VasX are indicated by curves marked with black stars. The last twenty residues of each VgrG2 are marked with red brackets on the diagram. **(C)** The interfaces between the different proteins are shown with the associated scores identified by AlphaBridge. **(D)** PAE (Predicted Aligned Error) plots generated by AlphaFold3 . The coloured bar corresponds to the predicted position errors (in Å). The red rectangles highlight the high-confidence regions (low error values) corresponding to the interfaces between VgrG2 (1) and VasX.
